# Environmental stress promotes entry into a pre-existing latent state in *Plasmodium falciparum*

**DOI:** 10.64898/2026.09.13.751295

**Authors:** Anush Aryal, Kwesi Forson, Nadia Prasad, Jennifer L. Guler

## Abstract

Artemisinin treatment eliminates most *Plasmodium falciparum* malaria parasites but leaves behind a small population capable of long-term survival and eventual recrudescence. Although these survivors have been associated with latency, it remains unclear whether drug treatment induces this state or selectively enriches parasites already predisposed for persistence. Environmental stress also enhances parasite survival following artemisinin exposure, suggesting that stress may promote persistence by increasing entry into latency. Here, we show that environmental stress promotes entry into latency and define the molecular features of this state using single-cell transcriptomics. Mild nutrient deprivation increased latency formation 2- to 3-fold following dihydroartemisinin (DHA) exposure, and latency frequency positively correlated with long-term parasite recovery. Single-cell RNA sequencing revealed a transcriptionally distinct early trophozoite population that emerged after DHA exposure and diverged from the normal developmental trajectory. A 200-gene classifier confirmed strong enrichment of this signature following artemisinin exposure across independent datasets. The classifier also detected rare latent-like parasites in untreated populations, suggesting the presence of a rare parasite subpopulation already predisposed for persistence. Consistent with previous studies, latent parasites exhibited reduced investment in growth-associated functions; however, our analyses revealed that latency is accompanied by extensive and coordinated transcriptional remodeling; they selectively maintained mitochondrial functions, redox homeostasis, nutrient acquisition pathways, lipid metabolism, and stress-responsive signaling. Latent parasites also preserved invasion functions and host-cell interactions, with many of the strongest latency markers encoding exported proteins associated with the erythrocyte and Maurer’s clefts. Environmental stress increased entry into this state without substantially altering its core transcriptional architecture, suggesting that stress influences the decision to enter latency more than the biology of latency itself. These findings support a model in which artemisinin survival arises, in part, through enrichment of a pre-existing latent subpopulation and identify promising candidates to target latency for future antimalarial interventions.

## Introduction

In 2024, nearly half of the global population lived in regions at risk for malaria, with an estimated 282 million cases and 610,000 deaths worldwide, most of which occurred in children under five years old (*World Malaria Report*, 2025). *Plasmodium falciparum* causes the most severe form of human malaria and accounts for the majority of malaria-related deaths. The widespread use of artemisinin-based combination therapy (ACT), the current first-line treatment for *P. falciparum* malaria, has substantially reduced disease burden (*World Malaria Report*, 2025). However, the parasite’s highly adaptive nature, coupled with the emergence and spread of artemisinin tolerance, continues to challenge malaria control efforts (Dondorp et al., 2009; Ariey et al., 2014; Lu et al., 2017). Maintaining progress towards malaria elimination will require both continued development of effective therapies and a deeper understanding of the mechanisms that enable parasite survival under stress.

As a parasite that alternates between the human host and the mosquito vector, *P. falciparum* encounters dramatic fluctuations in nutrient availability that require rapid metabolic adaptation (Saliba & Kirk, 2001). *P. falciparum* lacks a canonical mechanistic target of rapamycin (mTOR) pathway (Marreiros et al., 2023), a central regulator of nutrient sensing, metabolism, and growth (Sabatini, 2017; Saliba & Kirk, 2001). Instead, the parasite appears to rely on a set of conserved downstream stress-response and growth-regulatory mechanisms (Ariey et al., 2014; Babbitt et al., 2012; Kumar et al., 2004; Mbengue et al., 2015a; McLean & Jacobs-Lorena, 2017; Vaid et al., 2010; M. Zhang et al., 2017).

One aspect of stress adaptation is cross-tolerance, in which prior exposure to one stressor enhances survival of subsequent, distinct stressors (Mitchell et al., 2009; Pires et al., 2024). We previously demonstrated a form of cross-tolerance where mild nutrient stress increased *P. falciparum* survival following artemisinin treatment (Brown et al., 2023); depletion of a single nutrient (hypoxanthine or thiamine) was sufficient to trigger protective transcriptional reprogramming that prepared parasites to survive. These findings prompted studies to understand the mechanisms responsible for this adaptive shift, including the roles of nutrient sensing pathways and the potential induction of latency. This reversible growth-arrested state is generally characterized by reduced metabolic activity and enhanced drug tolerance (Babbitt et al., 2012; Teuscher et al., 2010a; Witkowski et al., 2010). In *P. falciparum*, latent parasites have been observed following exposure to artemisinin and related compounds, where a subpopulation of parasites survive treatment and resume growth (Teuscher et al., 2010a; Witkowski et al., 2010; M. Zhang et al., 2017). Despite increasing recognition of latency in malaria (Micchelli et al., 2024; Tripathi et al., 2024; Kiboi et al., 2025), the factors that promote entry into latency and the molecular features that define the latent state remain poorly defined.

A related unresolved question is whether latent parasites arise exclusively as a consequence of drug exposure or whether artemisinin treatment enriches a rare population already predisposed for persistence. Most studies have identified latent parasites only after drug treatment, making it difficult to distinguish between induction of latency and selective survival of pre-existing cells. Resolving this distinction is important because it fundamentally changes how parasite persistence is viewed: latency could represent either a transient stress response triggered by drug exposure or a conserved survival state that exists before treatment and is favored under drug pressure.

Determining whether nutrient deprivation influences latency induction may provide critical insight into how parasites survive drug pressure. To test this, we quantified latency in normal medium and nutrient deprivation conditions and detected a positive relationship between latency level and survival rates. Single-cell transcriptomic profiling of populations enriched for latent parasites revealed a distinct parasite population with conserved transcriptional reprogramming. Rather than undergoing widespread transcriptional shutdown, latent parasites remained transcriptionally active and selectively reorganized discrete programs while nutrient deprivation further reshaped the latent program. A 200-gene transcriptional signature detected latent-like parasite populations across independent single-cell datasets, including rare latent-like parasites in untreated populations, indicating that latency-associated features may pre-exist before drug exposure. These findings support a model in which environmental stress increases entry into a conserved latent state that contributes to parasite survival during antimalarial treatment.

## Results

### Nutrient deprivation increases parasite entry into latency following DHA exposure

To examine whether nutrient deprivation promotes parasite entry into latency, we applied an experimental framework adapted from prior work (Brown et al., 2023) and added phenotypic enrichment and analytical steps to directly quantify latency (**Fig. 1A**). We incubated asynchronous *P. falciparum* in low hypoxanthine (LH) medium or thiamine free (TF) medium, alongside normal medium (NM controls) (**Supplementary Table 1**; see *Methods*). We confirmed that nutrient deprivation was mild due to previously defined levels of growth reduction and minimal impact on mitochondrial membrane potential (Brown et al., 2023). Following nutrient deprivation, we exposed parallel cultures to dihydroartemisinin (DHA) in two forms; we treated cultures with 700nM DHA for 6 h to induce latency (Witkowski et al., 2013), and with 200nM DHA for 6 h to assess recovery and cumulative growth over 10 days (Brown et al., 2023) (**Fig. 1**).

**Figure 1.**
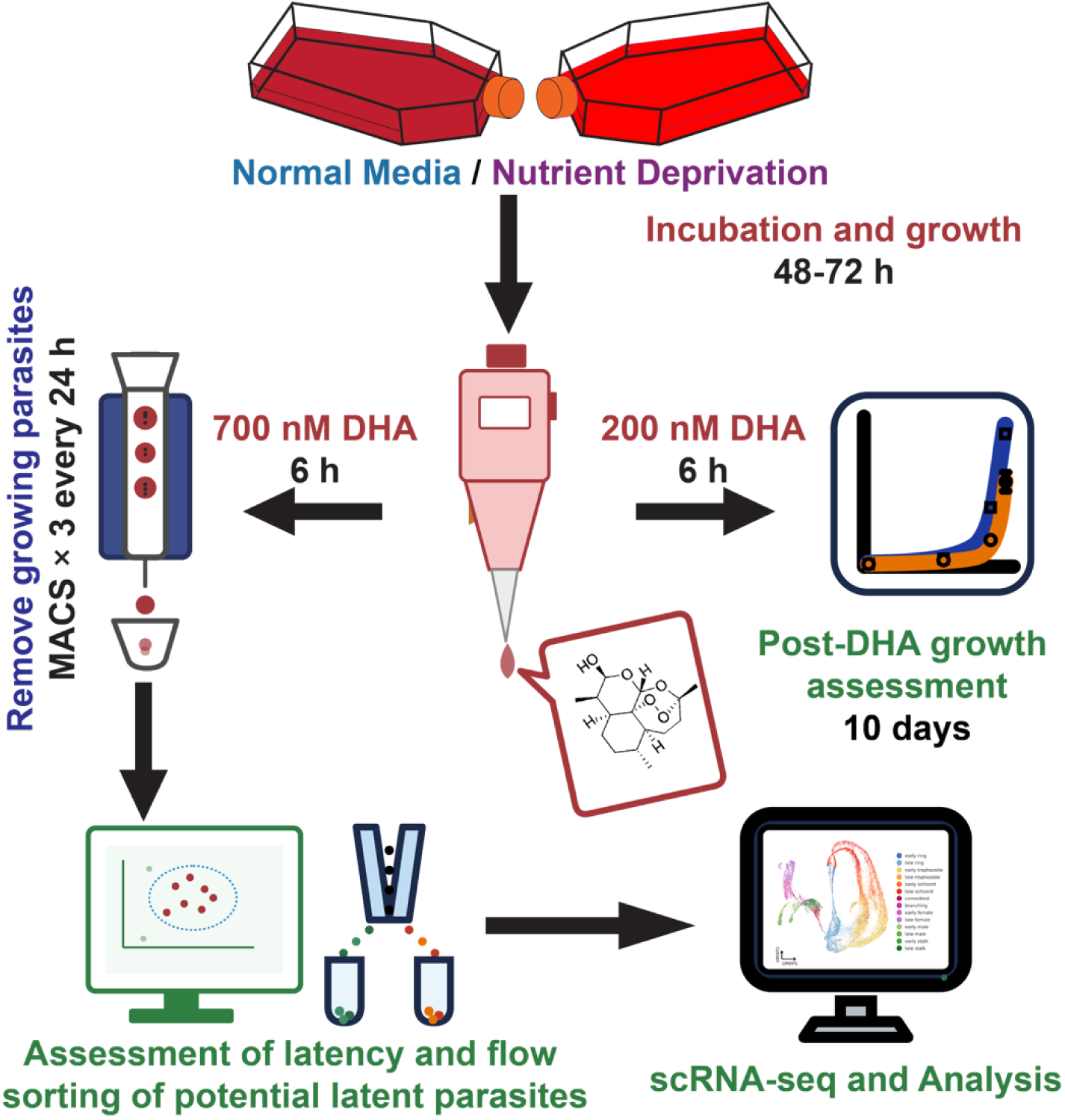
Experimental workflow. We cultured asynchronous *P. falciparum* under control (normal medium, NM) or nutrient-limiting (low-hypoxanthine, LH or thiamine-free, TF) conditions for 48-72h, then split cultures into two dihydroartemisinin (DHA) treatments. We treated one arm with 700nM DHA for 6h, depleted actively growing parasites by magnetic-activated cell sorting (MACS) once every 24h for three rounds, then quantified latency by flow cytometry and sorted potential latent parasites for single-cell RNA sequencing (scRNA-seq). We treated the second arm with 200nM DHA for 6h and monitored growth over 10 days to determine survival effects.

To remove actively growing parasites that survive DHA treatment and confound latency measurements, we performed standard depletion steps to enrich for latent parasites. For this step, we evaluated the impact of both D-sorbitol lysis (Lambros & Vanderberg, 1979a) and magnetic-activated cell sorting (MACS) (Ribaut et al., 2008) depletion methods. In practice, MACS preserved substantially more parasite material than D-sorbitol treatment after three successive rounds of actively growing parasite removal (D-sorbitol: 48.4% reduction in parasitemia, n = 3; MACS: 7.6% reduction in parasitemia, n = 6; P < 0.0001, unpaired t-test with Welch’s correction; **Supplementary Fig. 1**). Additionally, because D-sorbitol treatment relies on PSAC-mediated permeability in infected erythrocytes (Kutner et al., 1985; Pain et al., 2016) and nutrient stress increases PSAC mediated solute uptake (Potchen et al., 2025), this depletion method may lead to selective depletion of parasites with altered permeability characteristics (Nguitragool et al., 2011; Pillai et al., 2012). For these reasons, we chose to use MACS exclusively for subsequent enrichment steps. Similar to previous studies (Amaratunga et al., 2014; Brown et al., 2020; Connelly et al., 2021), we quantified latent parasites by flow cytometry based on parasite DNA content and preserved mitochondrial membrane potential (**Supplementary Fig. 2**).

Based on the criterion established in prior study (Brown et al., 2023), we restricted latency quantification to experiments that exhibited increased survival (≥100% day 10 difference, or ≥2-fold increase over NM controls) over 10 days following DHA treatment (**Supplementary Fig. 3A & 3B, Supplementary Table 2**). Under nutrient deprived conditions, parasites exhibited a ∼2- to 3-fold increase in latent parasites relative to NM controls (**Fig. 2A & B**). To examine the relationship between latency and long-term survival across experiments, we expanded the analysis to include all datasets where both latency measurements and day-10 growth data were available including those that did not pass the increased survival threshold. Across this combined dataset, fold change in latency showed a moderate association with fold change in day-10 survival (R² = 0.38, P=0.01, n = 17) (**Fig. 2C**). This association was similar within each nutrient deprivation condition (LH, R² =0.48; TF, R² =0.49).

**Figure 2.**
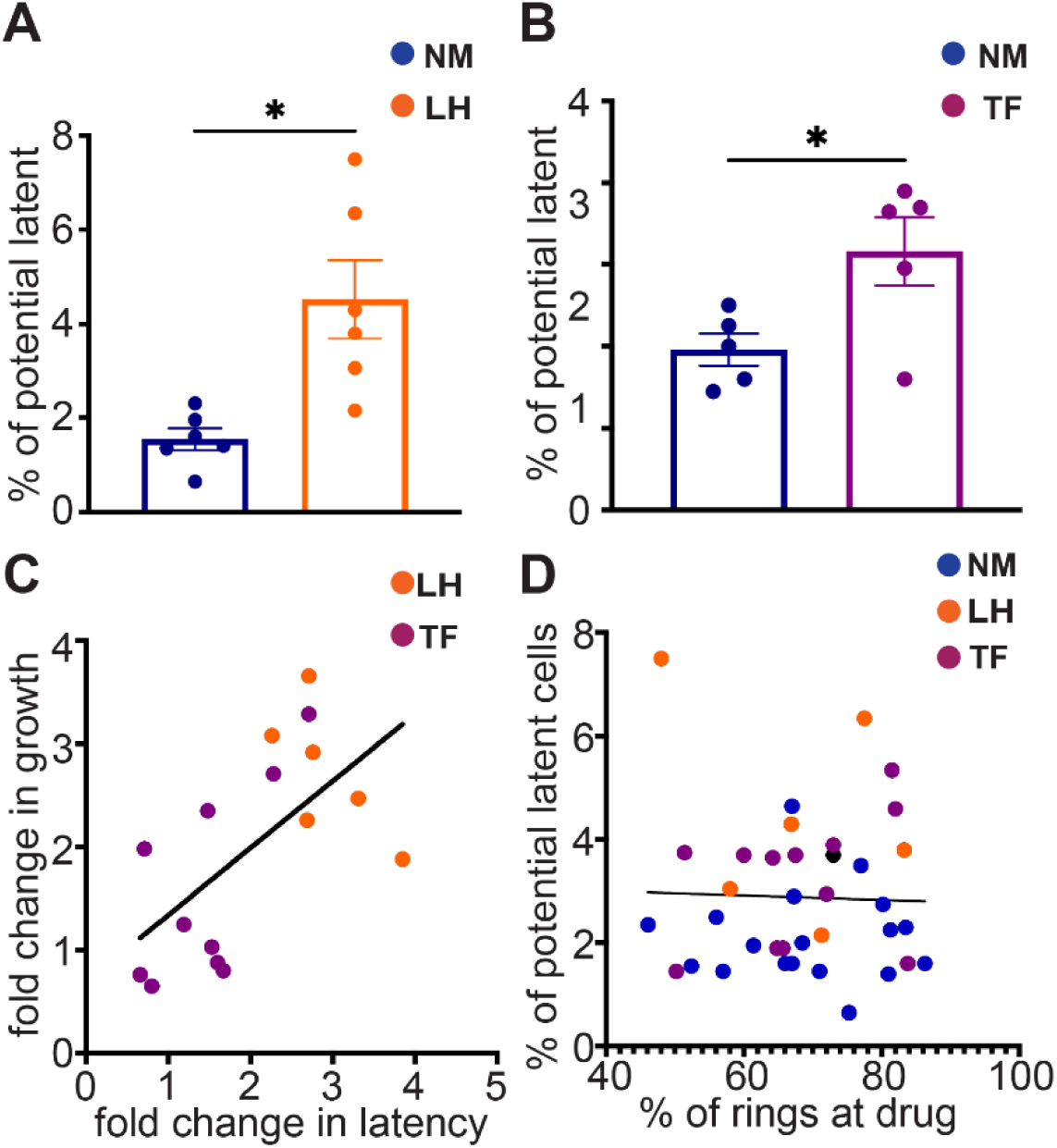
Nutrient deprivation increases the proportion of latent parasites following DHA treatment. [**A**] Percentage of potential latent parasites in low hypoxanthine (LH) medium compared to normal medium (NM), determined by SYBR Green- and MitoProbe-based flow cytometry. Bars show mean ± SEM with individual experiments overlaid. N = 6, *P < 0.05. [**B**] Percentage of potential latent parasites in thiamine-free (TF) medium compared to NM control, determined as in [A. n = 5, *P < 0.05. [**C**] Relationship between fold change in latency (flow cytometry, as in [**A**] and [**B**]) and growth measured by 10-day proliferation assays (R² = 0.38, n = 17). [**D**] Relationship between the percentage of ring-stage parasites at the time of dihydroartemisinin (DHA) treatment and the proportion of latent cells, expressed as the ratio of SYBR Green⁺/MitoProbe⁺ parasites to SYBR Green⁺ parasites. Each of the independent experiment contributes two points, one from NM and one from corresponding LH or TF (R² = 0.001, P=0.85, n = 36).

When we considered only experiments that had a ≥2-fold increase in both latency and day 10 survival, we observed no association between ring-stage abundance at the time of DHA treatment and latency (R² = 0.001, P=0.85, n = 36) (**Fig. 2D**). This result indicated that the presence of ring-stage parasites, which exhibit the highest DHA survival rates (Teuscher et al., 2010a; Witkowski et al., 2013), did not contribute to observed latency differences.

### Increased latency under nutrient deprivation occurs without eif2α activation

Based on prior studies implicating the eukaryotic initiation factor 2α (eIF2α) phosphorylation pathway in stress induced growth arrest (Babbitt et al., 2012) and artemisinin-induced latency (M. Zhang et al., 2017), we examined whether nutrient deprivation lead to elevated eIF2α phosphorylation in a bulk parasite population. We found that neither LH nor TF conditions increased eIF2α phosphorylation relative to controls (LH: ∼50% decrease, n = 5 per group, P = 0.95; TF: ∼8% decrease, n = 3 per group, P = 0.99; one-way ANOVA with Dunnett’s multiple comparisons, **Fig. 3A& 3B**). In contrast, DHA treated parasites displayed a marked increase in eIF2α phosphorylation (LH: ∼700% increase, n = 5, P = 0.014; TF: ∼3600% increase, n = 3, P < 0.0001), consistent with prior reports of drug-induced stress response (M. Zhang et al., 2017).

**Figure 3.**
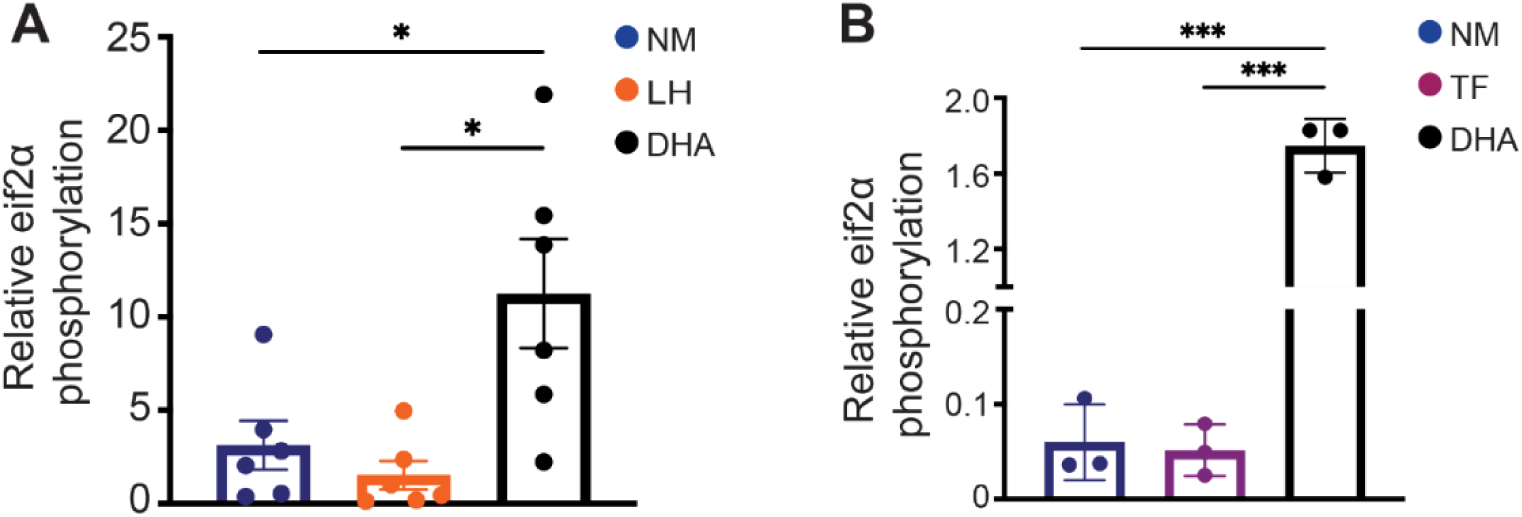
Nutrient deprivation does not cause a bulk increase in eIF2α phosphorylation. We calculated the relative abundance of phosphorylated eIF2α, calculated as the densitometric ratio of phosphorylated eIF2α /total eIF2α from Western blot analysis, in parasites cultured under **[A]** low hypoxanthine (LH, n=5) and **[B]** thiamine-free (TF, n=3) conditions compared to normal medium (NM) and dihydroartemisinin control (DHA, n=5 for panel A and n=3 for panel B). Bars show mean ± SEM with individual experiments overlaid.

### Single cell transcriptomics input is depleted of actively growing parasites but remains viable

To identify alternative pathways underlying entry into latency and to resolve transcriptional changes at the level of individual parasites, we performed single-cell RNA sequencing (scRNAseq) on parasites propagated under NM and LH conditions both before and after DHA exposure (pre- and post-DHA). As performed during latency quantification, we removed actively growing parasites after DHA treatment for 3 days prior to scRNAseq. For both conditions, we further enriched parasite populations using flow sorting that selectively retained viable infected erythrocytes (SYBR Green+/Mitoprobe+) while excluding uninfected (SYBR-) and dead parasites (Mitoprobe-) (**Supplementary Fig. S4**). To ensure that sorting conditions were compatible with scRNA-seq workflows, we evaluated multiple collection buffers and processing durations and monitored parasite viability for up to 6 h post-sorting, confirming stable viability under conditions used for cell isolation (**Supplementary Table 3**). Additionally, we confirmed that enriched populations consisted of viable latent parasites as opposed to dead cells resulting from DHA treatment (**Supplementary Fig. S4**). Sequencing metrics demonstrated robust data quality across all scRNAseq libraries, with recovery of 5,619–12,138 cells per sample, median detection of 463–765 genes per cell, and high fractions of reads mapping to cells (84.5–95.6%) (**Supplementary Table 4**).

### Reference-based stage annotation enables identification of a distinct latent population amongst early trophozoites

To assess the transcriptome of captured parasites, we placed individual cells within a staged developmental framework using the *P. falciparum* Malaria Cell Atlas (MCA) as a reference (Dogga et al., 2024) (**Supplementary Fig. 5A & 5B**). Reference mapping of combined post-DHA transcriptomes recovered the full intraerythrocytic developmental cycle and resolved the 12 developmental stages (**Supplementary Fig. 5B**), including early and late ring, early and late trophozoite, early and late schizont, sexually committed parasites, and early female gametocyte populations. This high-resolution staging approach established a consistent developmental framework across both NM and LH post-DHA conditions and revealed a divergent early trophozoite population (**Fig. 4A & 4B**), suggesting successful enrichment of latent parasites in post-DHA conditions.

**Figure 4.**
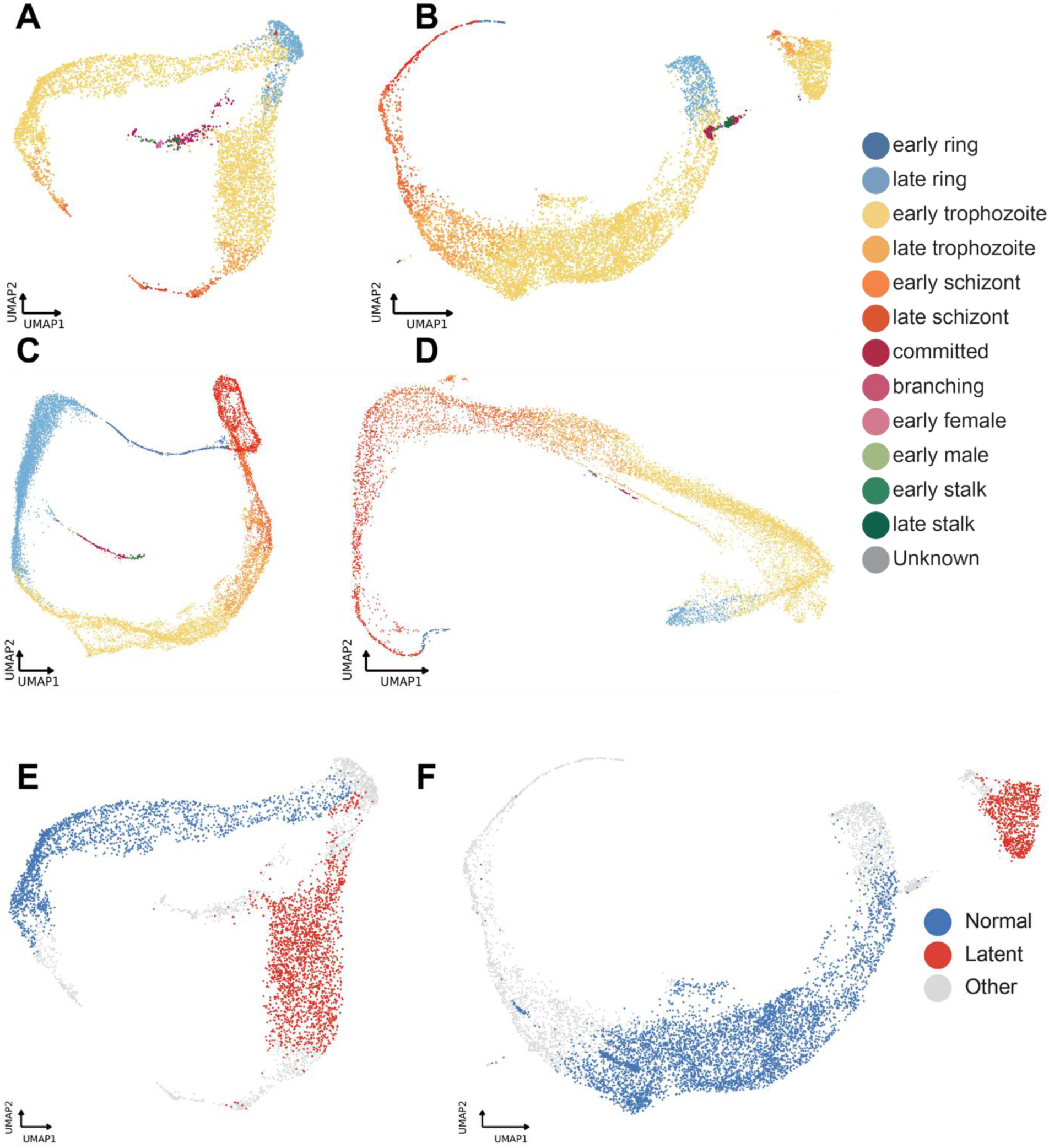
Reference mapping identifies a distinct parasite population enriched in post-DHA samples. UMAP representation of single parasites enriched for latency following DHA treatment in normal medium (NM) [**A**] and low-hypoxanthine (LH) conditions, colored by developmental stage. We assigned stages by projecting each cell onto the *P. falciparum* Malaria Cell Atlas reference and transferring the label of its nearest reference neighbors UMAP representation of untreated parasites cultured in NM [**C**] and LH [**D**] conditions, colored by developmental stage. [**E**] Co-embedded UMAP of post-DHA, latency-enriched parasites and untreated parasites from NM conditions ([**A**] and [**C**]), colored by classification as Normal early trophozoites (blue), Latent early trophozoites (red), or Other (grey). [**F**] Co-embedded UMAP of post-DHA, latency-enriched parasites and untreated parasites from LH conditions ([**B**] and [**D**]), colored by classification as Normal early trophozoites, Latent early trophozoites, or Other.

To validate parasite stage assignments in the post-DHA population, we examined expression of 15 curated marker genes across annotated populations and confirmed stage-specific patterns (**Supplementary Table 5, Supplementary Fig. 5C**). Ring, trophozoite, schizont, and sexual stage markers displayed conserved, stage-appropriate expression patterns consistent with known *P. falciparum* development. Trophozoites exhibited uniformly low expression of non-trophozoite markers, confirming that the divergence occurred within a bona fide early trophozoite context rather than through altered developmental timing. Within this validated developmental framework, post-DHA populations from both NM and LH conditions displayed a predominantly trophozoite-centered distribution with a clear bifurcation of the early trophozoite region (**Fig. 4A & 4B**).

To determine whether this divergence represented normal developmental variability or a distinct transcriptional state, we compared the post-DHA pattern to pre-DHA baseline populations. Pre-DHA parasites cultured in NM and LH conditions revealed a single trophozoite trajectory without evidence of branching (**Fig. 4C & 4D**). When we co-embedded cells from pre- and post-DHA conditions, one trophozoite trajectory was shared between both pre-DHA and post-DHA samples (blue population), whereas the divergent branch appeared exclusively in post-DHA parasites (red population, **Fig. 4E & 4F**). We defined this novel, post-DHA-specific branch as latent parasites and the shared trophozoite branch as normal stage-matched parasites.

To further confirm that the latent population represented an altered parasite state, we performed diagnostic scoring of cells within the post-DHA NM and LH datasets (**Table 1**). Latent cells retained high early trophozoite annotation confidence, with early trophozoite remaining their nearest classifier identity. However, they weakly expressed the normal early trophozoite transcriptional program; only 15.5% of NM latent cells and 6.4% of LH latent cells scored high for the normal early trophozoite signature, compared with 90.2% and 81.3% of normal early trophozoites, respectively. Few latent cells exceeded background for any alternative-stage signature, and sexual-stage marker expression remained at background levels. These findings indicate that latent parasites occupy a distinct early trophozoite-like state rather than representing a shift to an alternative developmental stage.

**Table 1.**
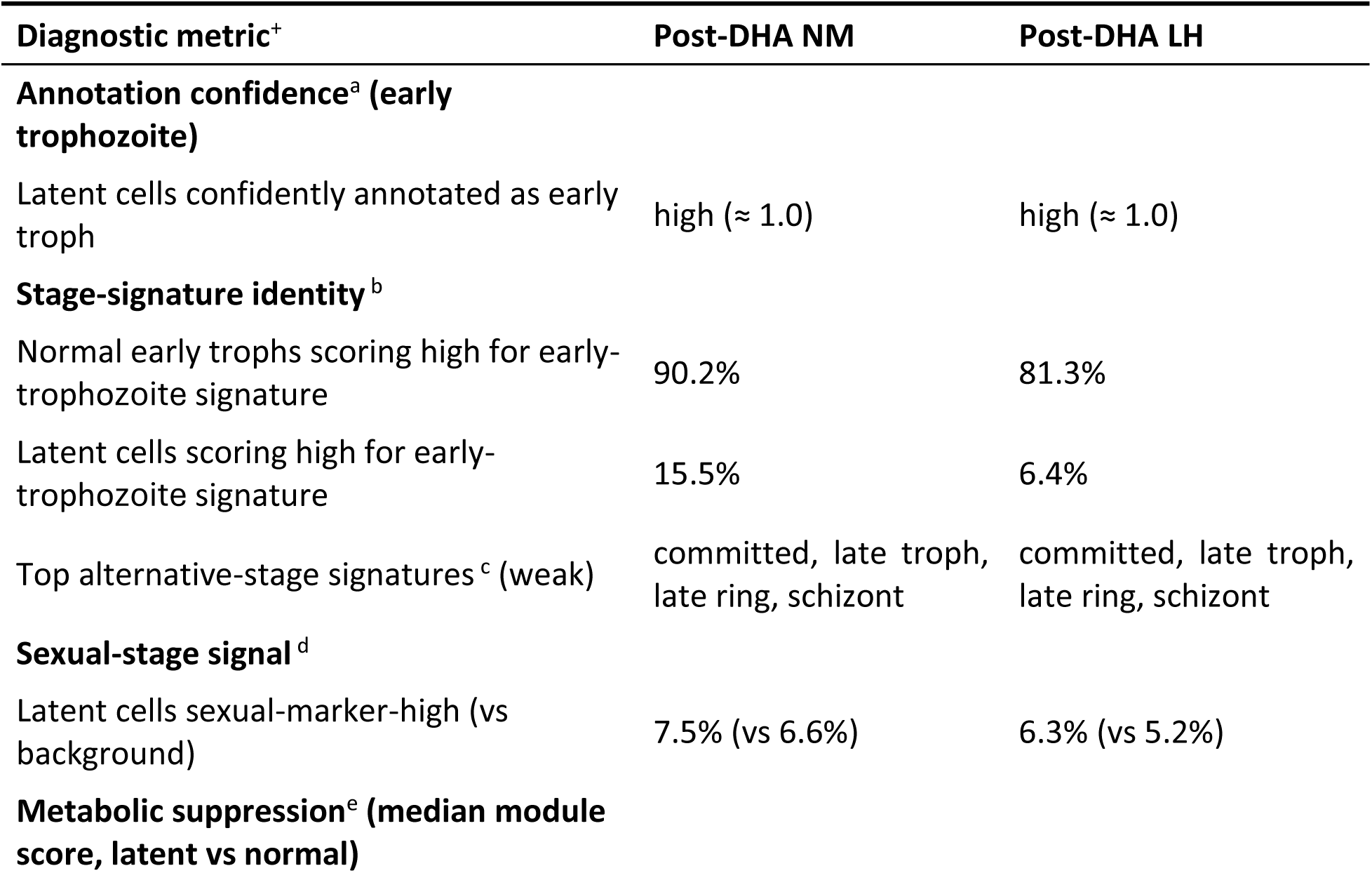
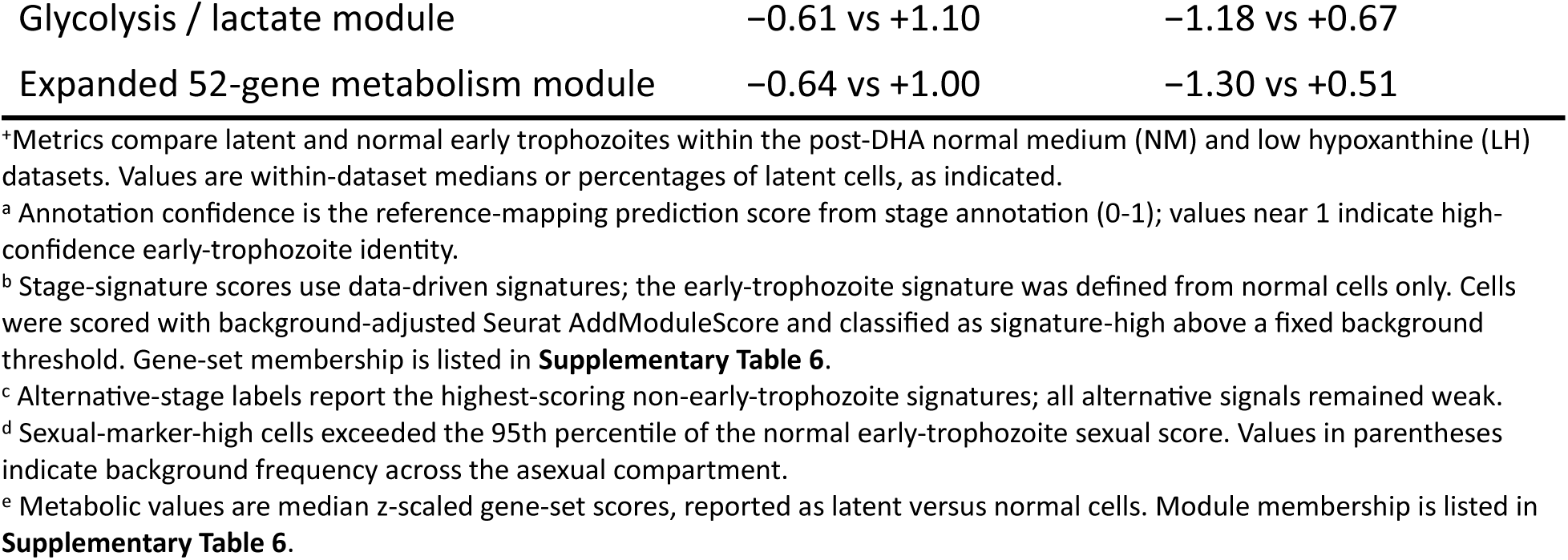
Evaluation of latent parasites compared to stage-matched early trophozoites.

| Diagnostic metric<sup>+</sup> | Post-DHA NM | Post-DHA LH |
| --- | --- | --- |
| **Annotation confidence<sup>a</sup> (early trophozoite)** |  |  |
| Latent cells confidently annotated as early troph | high ($\approx 1.0$) | high ($\approx 1.0$) |
| **Stage-signature identity<sup>b</sup>** |  |  |
| Normal early trophs scoring high for early-trophozoite signature | 90.2% | 81.3% |
| Latent cells scoring high for early-trophozoite signature | 15.5% | 6.4% |
| Top alternative-stage signatures<sup>c</sup> (weak) | committed, late troph, late ring, schizont | committed, late troph, late ring, schizont |
| **Sexual-stage signal<sup>d</sup>** |  |  |
| Latent cells sexual-marker-high (vs background) | 7.5% (vs 6.6%) | 6.3% (vs 5.2%) |
| **Metabolic suppression<sup>e</sup> (median module score, latent vs normal)** |  |  |
| Glycolysis / lactate module | -0.61 vs +1.10 | -1.18 vs +0.67 |
| Expanded 52-gene metabolism module | -0.64 vs +1.00 | -1.30 vs +0.51 |

\*Metrics compare latent and normal early trophozoites within the post-DHA normal medium (NM) and low hypoxanthine (LH) datasets. Values are within-dataset medians or percentages of latent cells, as indicated.
<sup>a</sup> Annotation confidence is the reference-mapping prediction score from stage annotation (0-1); values near 1 indicate high-confidence early-trophozoite identity.
<sup>b</sup> Stage-signature scores use data-driven signatures; the early-trophozoite signature was defined from normal cells only. Cells were scored with background-adjusted Seurat AddModuleScore and classified as signature-high above a fixed background threshold. Gene-set membership is listed in **Supplementary Table 6**.
<sup>c</sup> Alternative-stage labels report the highest-scoring non-early-trophozoite signatures; all alternative signals remained weak.
<sup>d</sup> Sexual-marker-high cells exceeded the 95th percentile of the normal early-trophozoite sexual score. Values in parentheses indicate background frequency across the asexual compartment.
<sup>e</sup> Metabolic values are median z-scaled gene-set scores, reported as latent versus normal cells. Module membership is listed in **Supplementary Table 6**.

To probe basic differences between latent and normal stage-matched parasites, we performed transcriptional metabolic module analysis that assesses relative activity of major metabolic pathways between parasite states. Across both post-DHA datasets and two independent module-scoring approaches, latent trophozoites showed reduced metabolic module activity relative to normal early trophozoites (**Table 1**). This suppression was selective rather than global (**Supplementary Fig. 8**), showing the strongest reductions in glycolysis/lactate metabolism, glutathione/redox metabolism, and pyridoxal phosphate synthesis. The TCA cycle module showed the least reduction across both conditions. Together, this analysis confirmed that latent parasites generally downregulate their metabolism compared to matched stages, but the selective remodeling warranted further investigation.

### Latency is associated with extensive transcriptional reprogramming

To characterize transcriptional differences between latent and stage-matched parasites, we performed differential expression (DE) analysis using Model-based Analysis of Single-cell Transcriptomics (MAST) (all genes shown in **Supplementary Table 7**). Since we identified latent parasites from both NM and LH conditions post-DHA treatment (**Fig. 4E & 4F**), we combined these datasets to define the core transcriptional features of latency independent of nutrient status. When we restricted the analysis to a high confidence DE gene set (adjusted P < 0.05, |log₂FC| ≥ 1), we detected 1,044 significantly upregulated genes and 1,116 significantly downregulated genes. This nearly equal shift of increased and decreased gene expression indicates that latency involves broad transcriptional remodeling (**Fig. 5A**, **Supplementary Tables 8 & 9**).

**Figure 5.**
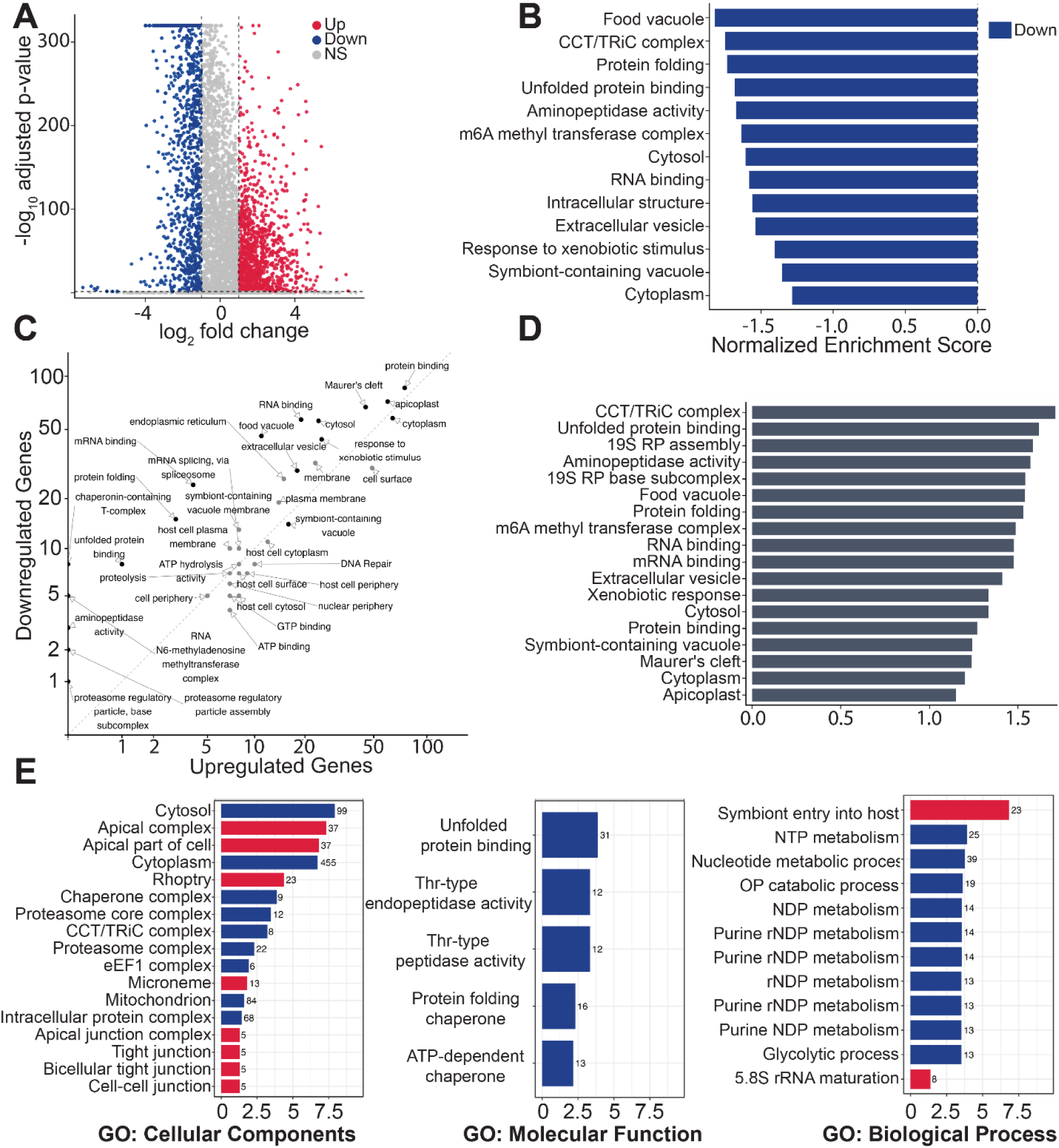
Latent parasites undergo selective transcriptional remodeling characterized by mixed-direction pathway regulation. Red, significantly upregulated; blue, significantly downregulated [**A**] Differentially expressed genes between latent and normal parasites (adjusted P < 0.05, |log₂FC| ≥ 1). Log₂ fold change was calculated using MAST, comparing latent parasites to stage-matched normal parasites in combined LH & NM conditions. [**B**] Gene Set Enrichment Analysis (GSEA) of pathways enriched in latent parasites relative to normal stage-matched parasites, restricted to pathways containing >5 genes. Pathways considered significantly enriched at adjusted P < 0.05. [**C**] Gene-level contribution analysis: the proportion of significantly upregulated, significantly downregulated, and non-significant genes within each tested pathway (using the same thresholds as in [**A**]). [**D**] Absolute GSEA analysis of the same pathways as in [**B**], ranking pathways by enrichment magnitude irrespective of direction. [**E**] Gene Ontology (GO) enrichment analysis across Biological Process, Molecular Function, and Cellular Component categories.

To identify coordinated functional programs within this DE gene set, we performed gene set enrichment analysis (GSEA) across 189 annotated pathways. Only 13 pathways reached the statistical significance (adjusted P < 0.05), and notably, they all reflected downregulation (**Fig. 5B, Supplementary Table 10**). Because standard GSEA relies on signed rankings (+ or -), pathways with a genuine mix of strongly upregulated and downregulated genes (**Fig. 5C**) may cancel out and remain undetected. Therefore, we ran GSEA on the absolute value of rankings (**Fig. 5D**) and identified 18 significantly enriched pathways. Conserved top hits across the standard and absolute GSEA included food vacuole, cytosol, protein binding, mRNA binding, cytosol, response to xenobiotic stimulus, extracellular vesicle, T-complex, and protein folding processes (**Fig. 5B & 5D**, **Supplementary Table 11**); however, absolute GSEA results highlighted new categories such as Mauer’s cleft and apicoplast pathways. When we assessed the proportion of genes altered in each pathway, T-complex, unfolded protein binding, protein folding, mRNA binding, RNA binding, and food vacuole processes skewed towards downregulation (75%-100% genes downregulated) while cytoplasm, protein binding, apicoplast, and symbiont-containing vacuole pathways had a nearly equal mix of upregulated and downregulated genes (**Supplementary Table 12**).

To understand more about altered transcription during latency, we performed Gene Ontology (GO) term enrichment analysis on the significantly altered gene sets across three GO domains (biological processes, BP, cellular components, CC, and molecular functions, MF). We identified 70 enriched GO terms overall: 60 downregulated terms and 10 upregulated terms (**Fig. 5E, Supplementary Table 13**). Reduced processes primarily involved in cytoplasmic metabolism and biosynthesis, specifically carbohydrate pathways (glycolysis GO term enrichment including 13 genes, carbohydrate derivative metabolic process: 49 genes), nucleotide process (nucleotide metabolic processes: 39 genes, purine nucleoside catabolism: 14 genes, pyridine nucleotide catabolism: 13 genes, nucleoside triphosphate metabolic process: 25 genes), energy derivation by oxidation of organic compounds (27 genes) and cellular respiration (26 genes), alongside broader metabolic and organellar terms (small molecule metabolic process: 77 genes, mitochondrion: 84 genes). Latent parasites also downregulated categories associated with protein quality control and turnover, including protein folding (36 genes), unfolded protein binding (31 genes), protein folding chaperones (16 genes), and the proteasome complex (22 genes). In contrast, upregulated GO terms centered around host-interaction and parasite export/invasion-associated programs, including the apical complex (37 genes), rhoptry (23 genes), symbiont entry into host (23 genes) microneme (13 genes), apical cellular structures (5 genes), as well as selected RNA maturation processes (8 genes).

Overall, GSEA and GO term analyses converged on several conserved features of latency, highlighting extensive remodeling of metabolism, RNA processing and post-transcriptional regulation, protein homeostasis, and cellular stress-response pathways, alongside changes in parasite-host interface functions. These recurring themes provided a framework for the following results sections.

### Latent parasites selectively redistribute metabolic activities across compartments

Due to strong metabolic suppression in latent parasites (**Table 1**, **Fig. 5B-E**), we specifically evaluated expression of metabolic genes in latent versus stage-matched normal. Overall, we found that latent parasites exhibited metabolic reprogramming rather than global suppression (**Fig. 6**), with distinct patterns of reduced, mixed, and elevated activity across subcellular compartments and metabolic pathways. As seen during GSEA, reduced processes primarily involved glycolysis and nucleotide synthesis and salvage, with nearly all significantly altered genes being downregulated. Latent parasites also strongly reduced hemoglobin digestion and proteolysis processes within the food vacuole with significant transcriptional downregulation of all falcipain and plasmepsin homologs (**Fig. 6**, average Log_2_FC of −1.2, **Supplementary Table 7 & 9**). However, latent parasites elevated transcription of the kelch 13 (*K13*)-associated endocytosis complex mediating hemoglobin uptake (*K13*, Log_2_FC of +1.5, and K13-interacting candidates *KIC1-KIC10*, average Log_2_FC of +1.5, **Supplementary Table 7 & 8**), indicating that hemoglobin uptake machinery remains transcriptionally supported despite suppression of downstream proteolytic digestion.

**Figure 6.**
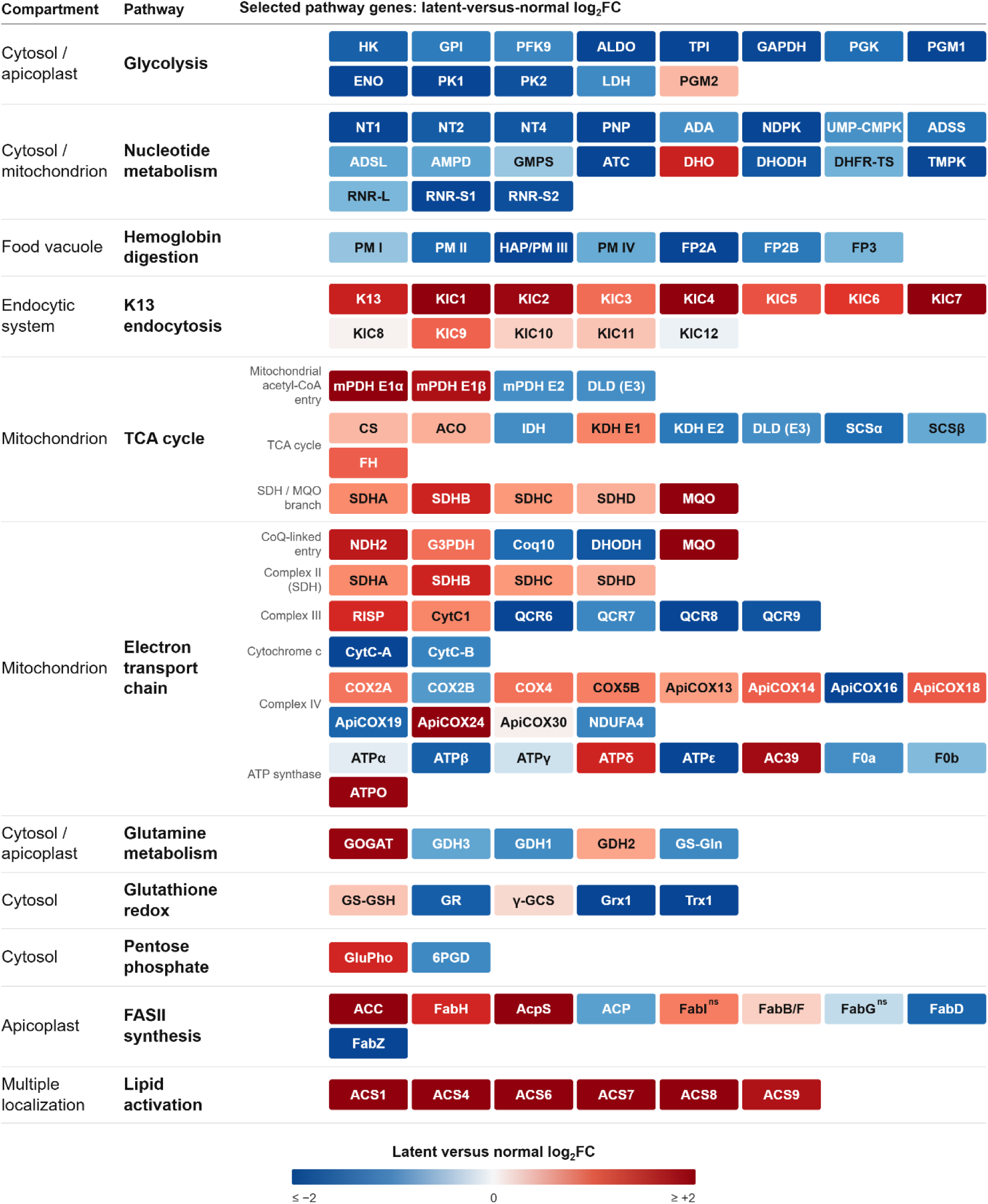
Metabolic reprogramming in latent *Plasmodium falciparum* parasites. Log₂ fold change (FC) in transcript abundance in latent versus stage-matched parasites for selected metabolic pathways. Each tile is one gene; blue indicates reduced and red increased abundance in latent parasites (see scale bar). Genes are grouped by pathway and listed under the compartment of the encoded enzyme. All changes are significant (p<0.05) unless marked “ns”. Gene identifiers follow PlasmoDB nomenclature; full PF3D7 accessions and log₂FC values listed in **Supplementary Table 14**: HK, hexokinase; GPI, glucose-6-phosphate isomerase; PFK9, phosphofructokinase 9; ALDO, fructose-bisphosphate aldolase; TPI, triosephosphate isomerase; GAPDH, glyceraldehyde-3-phosphate dehydrogenase; PGK, phosphoglycerate kinase; PGM1, phosphoglycerate mutase 1; ENO, enolase; PK1/PK2, pyruvate kinase 1 and 2; LDH, lactate dehydrogenase; PGM2, phosphoglucomutase 2; NT1/NT2/NT4, nucleoside transporters; PNP, purine nucleoside phosphorylase; ADA, adenosine deaminase; NDPK, nucleoside diphosphate kinase; UMP-CMPK, UMP-CMP kinase; ADSS, adenylosuccinate synthetase; ADSL, adenylosuccinate lyase; AMPD, AMP deaminase; GMPS, GMP synthetase; ATC, aspartate transcarbamoylase; DHO, dihydroorotase; DHODH, dihydroorotate dehydrogenase; DHFR-TS, dihydrofolate reductase– thymidylate synthase; TMPK, thymidylate kinase; RNR-L/RNR-S1/RNR-S2, ribonucleotide reductase large and small subunits; PM I–PM IV, plasmepsins I–IV; HAP, histo-aspartic protease (plasmepsin III); FP2A/FP2B/FP3, falcipains 2A, 2B and 3; K13, Kelch13; KIC1–KIC12, Kelch13-interacting candidates 1–12; mPDH, mitochondrial pyruvate dehydrogenase complex (also annotated as branched-chain ketoacid dehydrogenase, BCKDH), subunits E1α, E1β and E2; DLD, dihydrolipoamide dehydrogenase (E3 subunit shared with KDH); CS, citrate synthase; ACO, aconitase; IDH, isocitrate dehydrogenase; KDH E1/E2, α-ketoglutarate dehydrogenase subunits E1 and E2; SCSα/SCSβ, succinyl-CoA synthetase α and β subunits; FH, fumarate hydratase; SDHA–SDHD, succinate dehydrogenase subunits A–D; MQO, malate:quinone oxidoreductase; NDH2, type II NADH dehydrogenase; G3PDH, glycerol-3-phosphate dehydrogenase; Coq10, coenzyme Q-binding protein Coq10; RISP, Rieske iron–sulfur protein; CytC1, cytochrome *c*₁; QCR6–QCR9, ubiquinol–cytochrome *c* reductase subunits 6–9; CytC-A/CytC-B, cytochrome *c* isoforms; COX2A, COX2B, COX4 and COX5B, cytochrome *c* oxidase subunits; ApiCOX13–ApiCOX30, apicomplexan-specific cytochrome *c* oxidase subunits; NDUFA4, NADH dehydrogenase 1α subcomplex subunit 4 homolog; ATPα–ATPε, F₁ ATP synthase α, β, γ, δ and ε subunits; AC39, F₀ ATP synthase subunit d; F0a/F0b, F₀ ATP synthase subunits a and b; ATPO, oligomycin sensitivity-conferring protein; GOGAT, glutamate synthase; GDH1–GDH3, glutamate dehydrogenases 1–3; GS-Gln, glutamine synthetase; GS-GSH, glutathione synthetase; GR, glutathione reductase; γ-GCS, γ-glutamylcysteine synthetase; Grx1, glutaredoxin 1; Trx1, thioredoxin 1; GluPho, glucose-6-phosphate dehydrogenase–6-phosphogluconolactonase; 6PGD, 6-phosphogluconate dehydrogenase; ACC, acetyl-CoA carboxylase; AcpS, holo-acyl carrier protein synthase; ACP, acyl carrier protein; FabB/F, FabD, FabG, FabH, FabI and FabZ, type II fatty-acid synthesis enzymes; ACS1–ACS9, acyl-CoA synthetases.

As observed in our prior analysis (**Supplementary Fig. 6**), latent parasites maintained transcription of the mitochondrial TCA cycle over other metabolic processes. When evaluated in more detail, we observed a mixed pattern of up- and downregulation of TCA cycle transcripts (**Fig. 6**). Selective upregulation of TCA cycle components included those that reduce ubiquinone for use by the electron transport chain (*SDHA & SDHB,* average Log_2_FC of +1.1; *MQO,* Log_2_FC of +2.1, **Supplementary Table 8**). When evaluating the electron transport chain, we found that latent parasites elevated transcription of two additional enzymes that contribute to the pool of reduced ubiquinone (*NDH2* and *G3PDH*, average Log_2_FC of +1.2, **Supplementary Table 8**). The coordinated upregulation of these four distinct ubiquinone-reducing enzymes suggests that maintenance of the reduced ubiquinone pool and mitochondrial redox balance is a conserved feature of the latent state. Notably, latent parasites strongly downregulated another ubiquinone-dependent enzyme that also supports de novo pyrimidine biosynthesis (*DHODH*, Log2FC of −1.7, **Supplementary Table 9**), despite maintaining multiple alternative electron entry points into the electron transport chain. This pattern contrasts with the prevailing model that the primary role of the blood-stage electron transport chain is to support DHODH-dependent nucleotide biosynthesis (Ke et al., 2011; Painter et al., 2007).

While latent parasites predominantly down regulated respiratory complex III accessory factors (*QCR6-9,* average Log_2_FC of −2.0, **Fig. 6, Supplementary Tables 7-9**), they upregulated catalytic components *RISP* and *CYTC1* (average Log_2_FC of +1.1), suggesting preservation of core electron-transfer functions despite reduced investment in assembly. Similarly, latent parasites remodeled respiratory complex IV consistent with potential maintenance of mitochondrial membrane potential; we observed upregulation of multiple core components (*COX2A*, *COX4*, and *COX5B*, average Log_2_FC of +0.8) alongside reduced expression of complex IV-associated proteins (*ApiCOX16*, *ApiCOX19*, and *COX2B*, average Log_2_FC of −1.5). For ATP synthase, latent parasites decrease transcription of central catalytic and proton translocation components (*ATP-alph*a, *ATP-beta*, *F0a*, and *F0b*, average Log_2_FC of −0.9) while increasing transcription of structural components (*ATPO*, *AC39*, *ATP-delta*, average Log_2_FC of +1.6). This differential regulation suggests maintenance of ATP synthase architecture while reducing investment in ATP-producing capacity.

Latent parasites also displayed coordinated regulation of glutamine/glutamate metabolism and downstream pathways involved in redox homeostasis (glutathione metabolism; **Fig. 6**, **Supplementary Tables 7-9**). We observed expression changes consistent with increased glutamate availability for glutathione biosynthesis from salvaged glutamine; latent parasites upregulated glutamate production through *GOGAT* (Log_2_FC of +1.9), while reducing expression of *GS-Gln* (Log2FC of −1.1), which converts glutamate back to glutamine. Latent parasites also increased expression of *GluPho* (Log_2_FC of +1.4), indicating enhanced capacity for NADPH production through the pentose phosphate pathway, and slightly elevated expression of the glutathione biosynthetic enzymes *gamma-GCS* and *GS-GSH* (average Log_2_FC of +0.3). In contrast, latent parasites strongly downregulated expression of enzymes involved in glutathione utilization and recycling (*GR*, *Grx1*, and *Trx1*, average Log_2_FC of −2.4), indicating that latent parasites invest in the synthesis and accumulation of glutathione pools rather than glutathione turnover.

Finally, we observed a strong enrichment of lipid metabolic processes in latent parasites including initiating steps of fatty acid synthesis (*ACC, FABH, ACPS,* average Log_2_FC of +2.2, **Fig. 6, Supplementary Tables 7 & 8**). Some of the strongest upregulated genes in the entire dataset are from the acyl-CoA synthetase family (*ACS1*, *ACS4*, *ACS6*, *ACS7*, *ACS8*, and *ACS9,* average Log_2_FC of +3.2), which activate fatty acids for parasite use. The uniform upregulation of multiple acyl-CoA synthetases suggests that latent parasites maintain an enhanced capacity for fatty-acid activation and lipid remodeling, potentially supporting membrane maintenance and long-term cellular integrity. Taken together, our analysis of metabolic pathways indicated that latent parasites invest in hemoglobin uptake, mitochondrial maintenance, redox homeostasis, and lipid remodeling instead of growth-associated pathways.

### Latent parasites reorganize RNA processing, protein homeostasis, and integrated stress responses

When comparing latent to stage-matched normal parasites, we observed a coordinated reprograming of gene expression spanning RNA processing, proteostasis, nutrient sensing, phosphoinositide signaling, and autophagy. Consistent with the pathway-level enrichments described above, latent parasites broadly reduced expression of genes involved in RNA binding, translation initiation, ribosome biogenesis, and post-transcriptional regulation (**Fig. 5B-E**, **Supplementary Tables 10-13**). Genes of interest from this category included the RNA-binding proteins *PPR1* (Log₂FC of −1.7, **Supplementary Tables 7 and 9**), an essential apicoplast RNA-binding protein that protects organellar transcripts from degradation and has been proposed as an antimalarial target (Hicks et al., 2019), and *RACK1* (Log₂FC of −2.4), a ribosome-associated scaffold protein required for *P. falciparum* intraerythrocytic proliferation that binds ribosomes and actively translating polysomes (Blomqvist et al., 2017; Erath & Djuranovic, 2022) (**Supplementary Table 6**).

Proteostasis pathways exhibited a more complex pattern. Latent parasites strongly reduced expression of the entire TRiC/CCT chaperonin complex (*TCP1, CCT2, CCT3, CCT4, CCT5, CCT6, CCT7,* and *CCT8*; average Log₂FC of −1.7; **Fig. 5B**, **Supplementary Tables 7 and 9**). Similarly, latent parasites reduced *CYP19B* (Log₂FC of −1.2), a peptidyl-prolyl isomerase associated with protein-folding processes and stress response signaling (Kucharski et al., 2023). Consistent with these findings, both GO and GSEA analyses identified downregulation of protein folding, chaperonin, unfolded-protein binding, and TRiC-associated pathways (**Fig. 5B-E**; **Supplementary Tables 10-13**). Additional reductions within Prefoldin-associated pathways further suggest diminished capacity for folding newly synthesized proteins (**Table 2**).

**Table 2:**
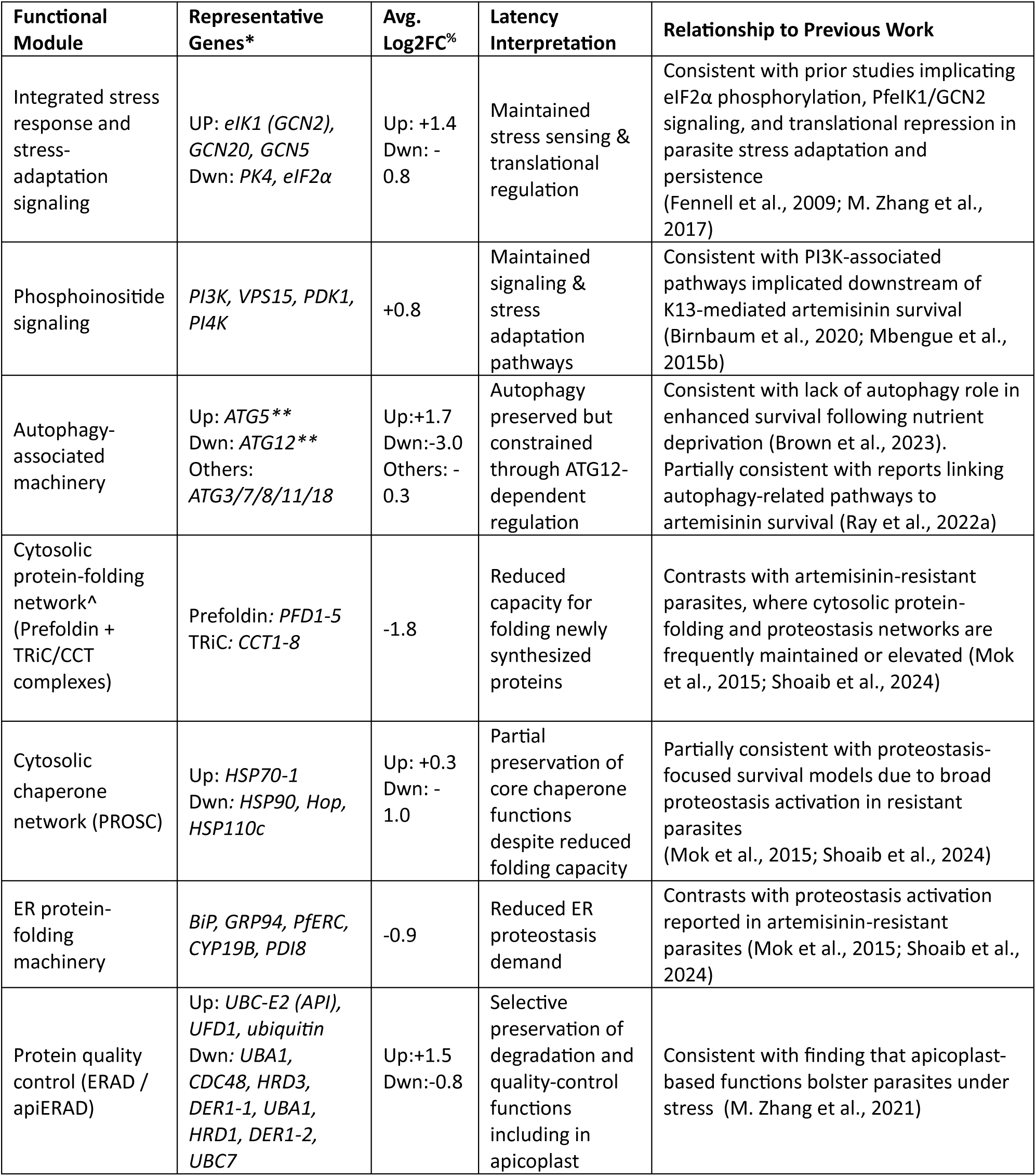
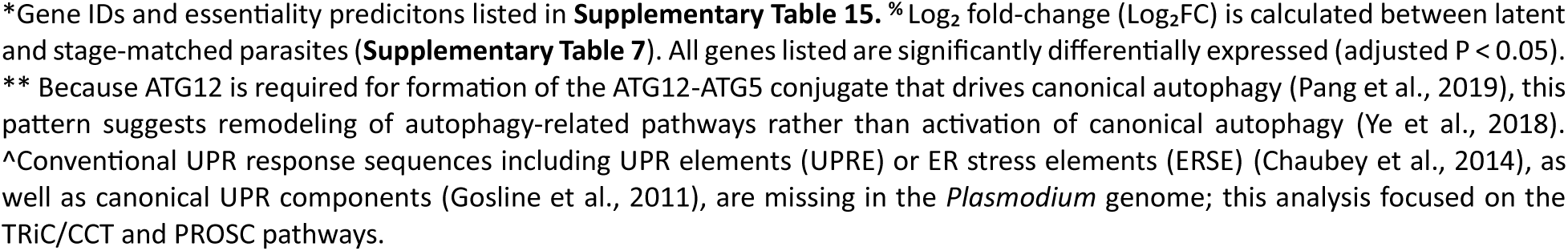
Significantly differentially expressed stress-response and proteostasis modules in latent parasites.

Despite prominent reductions, latent parasites selectively preserved protein quality-control pathways (**Table 2**). Enrichment analyses identified proteasome regulatory particle assembly, unfolded-protein binding, chaperonin complexes, and protein-folding processes among significantly altered pathways (**Fig. 5D**). At the pathway level, the ER-associated degradation (ERAD) pathway displayed mixed regulation, with increases of ubiquitin, one *UBC* allele, and *UFD1* (average Log₂FC of +1.5) alongside reductions of *HRD1, UBA1, CDC48, DER1-2,* a second *UBC* allele, and additional ERAD-associated components (average Log₂FC of −0.8, **Supplementary Tables 7-9**). The cytosolic PROSC chaperone network also exhibited mixed regulation (**Table 2**), indicating preservation of selected proteostasis functions. On the gene level, *GRP170*, an ER chaperone that interacts with the parasite BiP ortholog (Kudyba et al., 2019), showed minimal expression changes (Log₂FC of −0.1), consistent with retention of core quality-control functions within an overall reduced proteostasis network.

Latent parasites similarly remodeled stress-responsive pathways. Overlap between GO and GSEA analyses identified reduced expression across classical stress-response programs, including heat-stress, oxidative-stress, xenobiotic-response, and DNA-repair pathways (**Fig. 5B-E**; **Supplementary Tables 10-13**). However, targeted analysis revealed maintenance or modest activation of pathways previously implicated in artemisinin survival and stress adaptation (**Table 2**). Components of the eIF2α-mediated integrated stress response remained transcriptionally supported (**Table 2**); latent parasites significantly upregulated the GCN2 ortholog, *PfeIK1, GCN5,* and GCN20 (average Log₂FC of +1.4). Latent parasite also modestly increased components of phosphoinositide signaling, including *PI3K*, *VPS15*, *PI4K*, and *PDK1* (average Log₂FC of +0.8; **Supplementary Table 7 & 8**). Likewise, autophagy-associated genes displayed selective remodeling rather than uniform activation; we observed a strong upregulation of *ATG5* (Log₂FC of +1.7) alongside a dramatic reduction of its binding partner, ATG12 (Log₂FC of +3.0) (Pang et al., 2019), whereas most other autophagy-associated genes exhibited comparatively modest changes (**Table 2**).

Overall, the stress-response profile of latent parasites differs from both actively proliferating parasites and previously described artemisinin-resistant states (**Table 2**). Rather than broadly activating proteostasis and stress-survival pathways, latent parasites selectively preserve nutrient-sensing, phosphoinositide-signaling, autophagy-associated, and protein-quality-control functions while suppressing RNA processing, translation, and chaperone-mediated protein folding. Together, these findings support a model in which latent parasites prioritize cellular maintenance and regulatory control over growth-associated biosynthetic activity.

### Latency is associated with distinct variable surface antigen and trafficking profiles

Latent parasites increase pathways associated with parasite invasion and egress, including apical structures and rhoptries (**Fig. 5E**). To better characterize these transcriptional changes, we examined the expression of variable surface antigen and erythrocyte remodeling genes in latent versus stage-matched counterparts (**Fig. 7A, Supplementary Table 16**). While latent parasites downregulated the majority of PfEMP1 encoding *Var* genes across subfamilies (average Log_2_FC of −0.2, **Fig. 7B**), this reduction contrasted with the strong induction of other host-interaction and surface antigen gene families. These included serine repeat antigen (SERA, including *SERA1*, Log₂FC of +3.5, and *SERA5*, Log₂FC of +1.7), merozoite surface protein (including *MSP3*, *MSP7*, *MSP9*, and *MSP11*, all Log₂FC of >+3), and SURF family transcripts (including *SURF1.3*, log₂FC +5.2, with all-gene average Log_2_FC of +2.4 (**Fig. 7C**). Latent parasites also elevated transcription of additional invasion-associated ligands including *EBA175*, *EBA181*, *GAMA*, *RH1*, and *RH4* (**Supplementary Table 16**), indicating selective enhancement of host-interaction pathways despite broad suppression of PfEMP1-associated programs.

**Figure 7.**
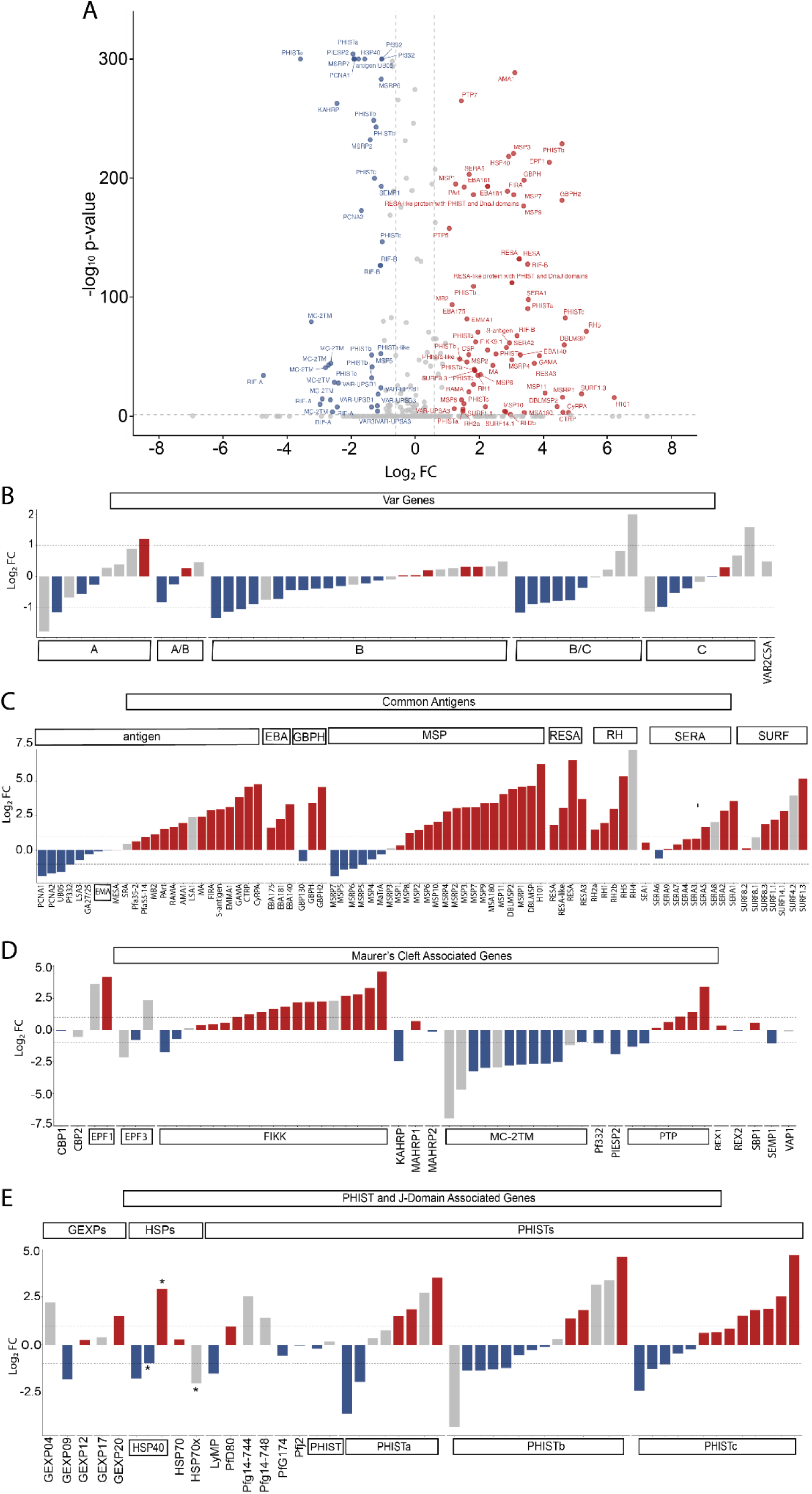
Latency is associated with selective remodeling of host-interaction and protein export genes. Red, significantly upregulated genes; blue, significantly downregulated genes, adjusted P < 0.05 and |log₂FC| ≥ 1; non-significant genes are shown in gray. Latent and stage-matched parasites are compared as pooled groups (NM and LH datasets). All genes listed in **Supplementary Table 16**. [**A**] Differential gene expression (log₂ FC, fold change) between transcriptionally defined latent and stage-matched parasites. Genes labeled with PlasmoDB gene annotations. Dotted lines indicate |log₂FC| = 1. **[B-E]** Differential gene expression from categories of exported proteins: [**B**] Var genes, [**C**] Maurer’s cleft-associated proteins, [**D**] J-Dot associated proteins and PHIST genes, [**E**] Surface antigen genes. *known components of Jdot complexes (Külzer et al., 2010; Anas et al., 2020).

Latent parasites broadly upregulated erythrocyte-remodeling transcripts including members of the FIKK-family kinases, which interact with the host cell cytoskeleton (Davies et al., 2020); *FIKK11* exhibited one of the largest increases in the entire dataset (Log₂FC of +5.1) and selectively marked latent parasites (see more details below, **Table 3**). Latent parasites also increased several additional FIKK family members, including *FIKK9.1*, *FIKK9.2*, *FIKK9.5*, and *FIKK9.6* (average Log₂FC of +2.5), while reducing only *FIKK10.1* and *FIKK10.2* (average Log₂FC of −1.2) (**Fig. 7D**). These patterns suggested selective activation of specific host-cell remodeling modules rather than generalized induction of the entire FIKK family.

**Table 3.** Latency-associated marker set showing dominance of proteins localized to the host cell.

| Gene ID | Annotation | Log<sub>2</sub>FC | % detection (latent/normal) | Localization<sup>a</sup> | Role / biological theme<sup>b</sup> |
| --- | --- | --- | --- | --- | --- |
| PF3D7_1149300 *** | FIKK11 | +5.1 | 11.9% / 0.9% | Host cell periphery | Protein kinase; host-cell perturbation |
| PF3D7_1301200 *** | GBPH2 | +5.1 | 13.0% / 1.9% | Host cell | Glycophorin-binding (by homology)<sup>c</sup> |
| PF3D7_0114000 *** | EPF1 | +4.8 | 17.9% / 2.6% | Maurer's cleft | Protein binding; exported remodeling |
| PF3D7_1000600 *** | RIFIN | +4.1 | 13.2% / 1.8% | Host cell plasma membrane | Antigenic variation |
| PF3D7_1401000 *** | GBPH | +4.0 | 15.6% / 4.8% | Host cell | Glycophorin-binding (by homology)<sup>c</sup> |
| PF3D7_0113700 *** | HSP40, type II | +3.5 | 24.5% / 6.5% | Maurer's cleft; host cell cytosol | Chaperone / unfolded-protein binding |
| PF3D7_0924700 *** | SF3A3 | +3.4 | 19.1% / 6.3% | Spliceosomal complex | mRNA splicing; RNA binding |
| PF3D7_1328500 *** | Exported lipase homolog 4 (XLH4<sup>d</sup>) | +3.0 | 23.8% / 9.8% | parasite cytoplasm<sup>d</sup> | Lipid scavenging |
| PF3D7_1201100 *** | RESA-like PHIST-DnaJ | +2.4 | 34.8% / 10.2% | Host cell cytoplasm | Exported PHIST/DnaJ remodeling<sup>e</sup> |

\*\*\* denotes adjusted $P < 10^{-10}$ . Log<sub>2</sub>FC is calculated from latent parasites/stage-matched normal parasites MAST analysis. % detection is the fraction of latent versus normal early trophozoites expressing each gene. <sup>a</sup>Localization from Gene Ontology cellular-component annotations (PlasmoDB release 68); “-”: no GO localization term. <sup>b</sup>Role/theme is summarized from GO molecular-function and biological-process terms where available. <sup>c</sup>GBPH/GBPH2 and the RESA-like PHIST-DnaJ protein lack informative GO function/process terms; their roles are inferred from gene-family annotation or homology. <sup>e</sup>Characterized by (Liu et al., 2024)

Latent parasites also altered transcripts associated with parasite-derived structures that sit in the erythrocyte cytoplasm, including Jdots and Maurer’s clefts (**Fig. 5B & 5D; Supplementary Table 16**). Exported type II *HSP40*, which is associated with Jdots (Almaazmi et al., 2022), selectively marked latent parasites (**Table 3**). The PHIST protein family also contributes to these trafficking structures (Oberli et al., 2016; Warncke et al., 2016; Almaazmi et al., 2022), and latent parasites displayed mixed regulation across PHIST subtypes (**Fig. 7E**); *PHISTa* family members included both some of the most strongly induced and most strongly repressed exported proteins, whereas *PHISTb* family members were predominantly downregulated and *PHISTc* family members were largely upregulated (**Supplementary Table 16**). Within Maurer’s cleft pathways, latent parasites strongly reduced transcription of *MC-2TM* family members*, KAHRP*, and *SEMP1* (average Log_2_FC of −2.8), while increasing transcripts contributing to Maurer’s cleft formation (*EPF1* and *FIKK9.1,* average Log_2_FC of +3.4) and PfEMP1 trafficking proteins (PTP: *PTP1*, *PTP7*, and *PTP5*, average Log_2_FC of +1.4, **Fig. 7D & 7E**). Together, these opposing patterns suggest that latent parasites retain and selectively remodel export machinery while reducing components associated with canonical virulence architecture and cytoadherence.

When we sought to identify a transcriptional signature of latency independent of pathway-level analyses, we identified nine strongly upregulated transcripts in latent parasites (Log_2_FC of between +2.4 to +5.1, **Table 3**). Consistent with selective activation within the latent population, a substantially larger fraction of latent parasites strongly expressed these marker genes compared to a minority of stage-matched parasites (**Table 3**). Strikingly, seven of the nine latency markers localize to the erythrocyte including the host-cell periphery, cytoplasm, or Maurer’s clefts, linking latent parasites directly to pathways involved in host-cell remodeling. Many of these proteins remain relatively uncharacterized but strongly upregulated including *FIKK11* (Log_2_FC of +5.1), *GBPH2* (+5.1), *GBPH* (+4.0), and *XLH4* (+3.0) (**Table 3**). Despite being a FIKK-family kinase, very little is known about FIKK11. GBPH and GBPH2 interact with the infected erythrocyte, where they have been reported to be co-regulated with protein-export machinery in trophozoites developing in sickle-trait erythrocytes (Saelens et al., 2021). XLH4 has only recently been implicated in the utilization of host lysophosphatidylcholine for fatty acid scavenging from the host environment (Liu et al., 2024). This strong enrichment of exported and host-cell-associated proteins suggests that remodeling of the infected erythrocyte is a defining feature of the latent transcriptional state.

### Annotation of previously uncharacterized genes reinforce translational, redox, and signaling capacity

A substantial fraction of the latency-associated transcriptional program included genes of unknown function, limiting our mechanistic interpretation. Of the differentially expressed genes between latent and normal parasites, 28% lacked any functional annotation and were labeled only as conserved unknown, hypothetical, or exported proteins of unknown function (**Supplementary Table 17**). To assign putative functions to these genes, we applied PlasmoFP, a structure-informed deep-learning function predictor trained on Gene Ontology (GO) relationships across the genetically related eukaryotic organism SAR supergroup (Srivastava et al., 2025). At an effective FDR ≤ 0.05, 406 of the uncharacterized latency genes (out of 1388 total) received at least one PlasmoFP GO prediction. Critically, 275 genes that previously carried no GO annotation of any kind acquired a predicted function, substantially reducing the annotation gap within the latency program (**Supplementary Table 17**). The predicted functions covered broad categories including binding, catalytic, transport, ribosomal, and RNA-processing themes, indicating that uncharacterized genes participated in the coordinated remodeling during latency (**Supplementary Table 17**). Among upregulated uncharacterized genes, we recovered two genes predicted to encode transmembrane transporters (**Table 4**). Additionally, PlasmoFP assigned 29 genes as structural components of the ribosome, 16 of which were upregulated in latent parasites and predicted to be essential. Other predictions extended the RNA-processing signature, including pseudouridine synthase, and nucleic acid catalytic activities. Finally, the uncharacterized gene list included two putative mitochondrial oxidoreductase genes and a set of kinase- and phosphatase-regulatory genes (**Table 4**), consistent with the preserved redox and signaling activity that distinguishes latency from a global metabolic shutdown.

**Table 4.** PlasmoFP-predicted functions among uncharacterized latency genes.

|  | Gene ID (sig) | PlasmoFP predicted function | Log<sub>2</sub>FC | Essential?<sup>a</sup> | Location<sup>b</sup> |
| --- | --- | --- | --- | --- | --- |
| *Transmembrane transport* | PF3D7_1307500 *** | transmembrane transporter | +1.15 | Essential | memb |
|  | PF3D7_1329200 *** | transmembrane transporter | +1.07 | Dispensable | — |
| *Ribosome translation* / | PF3D7_1122700 *** | structural constituent of ribosome | +4.23 | Essential | memb |
|  | PF3D7_1005900 *** | structural constituent of ribosome | +3.86 | Essential | apico |
|  | PF3D7_1224800 *** | structural constituent of ribosome | +2.96 | Essential | — |
|  | PF3D7_0521200 *** | structural constituent of ribosome | +2.37 | Essential | mito |
|  | PF3D7_0921500 ** | structural constituent of ribosome | +2.29 | Essential | memb |
|  | PF3D7_0806900 *** | structural constituent of ribosome | +2.22 | Essential | apico, memb |
|  | PF3D7_1306100 *** | structural constituent of ribosome | +2.04 | Essential | — |
|  | PF3D7_1460200 ** | structural constituent of ribosome | +1.96 | Essential | mito |
|  | PF3D7_0728300 *** | structural constituent of ribosome | +1.91 | Essential | mito |
|  | PF3D7_1343500 *** | structural constituent of ribosome | +1.91 | Essential | apico |
|  | PF3D7_0903000 *** | structural constituent of ribosome | +1.86 | Essential | mito, memb |
|  | PF3D7_1470300 *** | structural constituent of ribosome | +1.64 | Essential | — |
|  | PF3D7_0911600 *** | structural constituent of ribosome | +1.59 | Essential | apico, memb |
|  | PF3D7_0305400 *** | structural constituent of ribosome | +1.59 | Essential | RNP |
|  | PF3D7_1012100 *** | structural constituent of ribosome | +1.30 | Essential | mito |
|  | PF3D7_1217000 *** | structural constituent of ribosome | +1.29 | Essential | ribo, RNP |
| *RNA processing* | PF3D7_0915500 *** | pseudouridine synthase activity | +3.75 | Essential | — |
|  | PF3D7_1147600 *** | catalytic activity, acting on a nucleic acid | +2.33 | Dispensable | — |
| *Oxidoreductase* | PF3D7_0503900 *** | oxidoreductase activity | +2.99 | Dispensable | mito, memb |
|  | PF3D7_1232600 *** | oxidoreductase activity | +1.04 | Essential | mito |
| *Kinase phosphatase* / | PF3D7_1318900 *** | kinase regulator activity | +2.47 | Dispensable | — |
|  | PF3D7_0418900 *** | phosphatase activity | +2.82 | Essential | — |
|  | PF3D7_0313000 *** | binding | +1.53 | Dispensable | — |

\*, level of significance of differential expression (adjusted P): \*\*\* $P < 10^{-10}$ , \*\* $P < 10^{-4}$ , \* $P < 0.05$ . Log<sub>2</sub>FC is calculated from latent parasites/stage-matched normal parasites MAST analysis. Predicted functions are PlasmoFP assignments at effective FDR $\leq 0.05$ (Srivastava et al., 2025).<sup>a</sup>Essentiality from the piggyBac saturation-mutagenesis screen (Zhang et al., 2018): “Essential”: non-mutable in CDS, “Dispensable”: mutable. <sup>b</sup>Predicted subcellular location from Gene Ontology cellular-component annotations (PlasmoDB release 68); “—”: no localization term. Memb, membrane; Apico, apicoplast; Mito, mitochondrion; Ribo, ribosome; RNP, ribonucleoprotein complex. All genes listed in **Supplementary Table 17**.

### Prior nutrient deprivation reinforces a resource-conserving latent transcriptional program

We next investigated the transcriptional differences in latent parasites from NM and LH conditions and found that parasites from both media types largely preserved the core latency program (**Fig. 8A**, **Supplementary Table 18-20**). At high confidence (adjusted P < 0.05, |Log₂FC| ≥ 1), we detected relatively few differentially expressed genes between media conditions (24 upregulated and 61 downregulated genes), indicating that nutrient limitation does not fundamentally alter the latent transcriptional state. Consistent with this observation, conventional GSEA detected no significantly enriched pathways when genes were ranked by signed fold-change (**Supplementary Table 21**). In contrast, absolute GSEA revealed eight significantly enriched gene sets including heterochromatin, trans-Golgi network, ER membrane complex, exosome/RNase complex, mitochondrial respiratory chain complex IV, phosphatidylinositol-3-phosphate binding, plastic organization, and protein binding processes (**Fig. 8B, Supplementary Table 22**). GO term analysis similarly identified a limited set of altered categories, with translation and protein biosynthetic processes reduced and rhoptry- and apical complex-associated genes increased (**Fig. 8C**, **Supplementary Table 23**). These results suggest that prior nutrient deprivation remodels specific components of the latent program rather than establishing a distinct transcriptional state.

**Figure 8.**
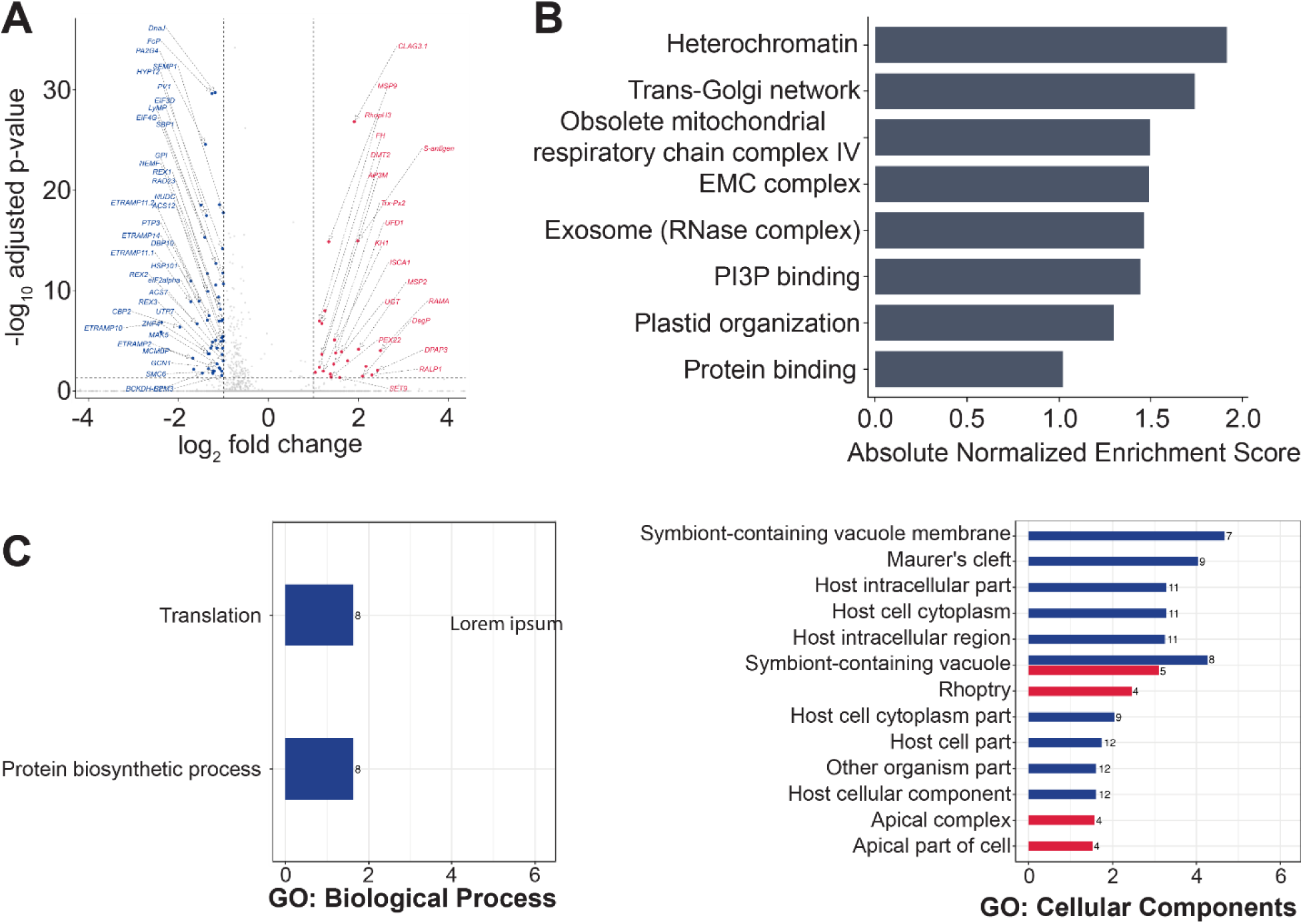
Nutrient limited condition has a limited but selective impact on the established latent transcriptional program. Red, significantly upregulated; blue, significantly downregulated. [**A**] Differentially expressed genes comparing latent parasites from low-hypoxanthine (LH) and normal (NM) medium (adjusted P < 0.05, |log₂FC| ≥ 1). [**B**] Absolute GSEA analysis of LH latent vs. NM latent parasites. [**C**] GO enrichment analysis. Differentially expressed genes between LH latent and NM latent parasites are enriched for translation and protein biosynthesis (Biological Process) and for host-associated cellular localization, including host cell cytoplasm, host intracellular region, symbiont-containing vacuole membrane, Maurer’s cleft, and rhoptry-associated components (Cellular Component).

Within this conserved program, LH latent further reduced the expression of pathways associated with growth and biosynthesis. Downregulated genes included multiple translation initiation and elongation factors (*eIF2α, eIF3D, eIF4G, EF1α, SUI1,* and *GCN1,* average Log_2_FC of −1.2; **Supplementary Table 18 & 20**) together with components of ribosome biogenesis and RNA maturation (*UTP7, MAK5, DBP10, NUDC, NUP116,* and a THUMP-domain protein, average Log_2_FC of −1.1), indicating additional restriction of protein synthesis capacity. LH latent parasites also reduced expression of parasitophorous vacuole and erythrocyte remodeling factors, including multiple ETRAMP family members (*ETRAMP11.2, ETRAMP2, ETRAMP11.1, ETRAMP14*, and *ETRAMP10,* average Log_2_FC of −1.8). Reductions also covered multiple exported-protein trafficking components, including the REX proteins (*REX1-3*; average log₂FC of −1.35), together with *MAHRP1, SBP1, SEMP1,* and *PTP3* (average Log₂FC of −1.2), consistent with reduced investment in Maurer’s cleft function and host-cell remodeling. For metabolism, LH latent parasites partially reversed components of the metabolic program observed during latency establishment, including the lipid-activation pathway (ACS7, ACS12, average Log_2_FC of −1.2) and mitochondrial carbon metabolism (*BCKDH-E2*, Log_2_FC of −1.5). These changes suggest that nutrient limitation drives a deeper resource-conserving latent state with reduced investment in protein production, host-cell remodeling, and metabolism.

In contrast, LH latent parasites selectively increased expression genes associated with growth recovery, nutrient acquisition, and organelle maintenance. Upregulated transcripts included multiple invasion- and egress-related factors (*RAMA, RALP1, RhopH3, DPAP3, DegP, MSP2,* and *MSP9,* average Log₂FC of +2.0; **Supplementary Table 18 & 19**), consistent with GO enrichment of rhoptry and apical-complex components. Notably, LH latent parasites increased expression of both *RhopH3* and *CLAG3.1* (average Log₂FC of +1.5), which together contribute to formation of the plasmodial surface anion channel (PSAC) (Ito et al., 2017; Nguitragool et al., 2011). This finding suggests that latent parasites maintain enhanced capacity for nutrient uptake despite broad suppression of biosynthetic pathways, even beyond the period of nutrient restriction.

LH latent parasites also increased expression of genes involved in trafficking and transport including *DMT2,* an essential apicoplast transporter (Sayers et al., 2018), as well as *AP3M* and *PEX22,* two uncharacterized proteins with predicted roles in membrane trafficking (Log₂FC of +1.2= +1.2), further supporting preservation of resource acquisition and intracellular trafficking functions. In parallel, transcriptional changes in mitochondrial genes pointed toward maintenance of organelle function. LH latent parasites elevated expression of *ISCA1*, a component of Fe-S cluster biogenesis (Mohammad Sadik et al., 2021) (Log₂FC of +1.6), and *FH* (Log₂FC of +1.2), the enzyme preceding MQO in the TCA cycle (Rajaram et al., 2022). The concurrent increase in *ISCA1* and *FH* alongside reduced expression of *BCKDH-E2* suggests selective preservation of mitochondrial integrity and electron-transfer capacity rather than increased mitochondrial carbon flux. Combined with enrichment of mitochondrial respiratory chain complex IV genes identified by absolute GSEA (**Fig. 8B**), these patterns are consistent with selective preservation of mitochondrial integrity and electron-transfer capacity during latency.

### Latent-like parasites occur at low basal frequencies but are strongly enriched following artemisinin treatment

We next sought to construct a classifier that could distinguish latent from normal parasites to determine the frequency of latency in our current study as well as independent single-cell datasets. After evaluating more than 150,000 combinations of gene panels, scoring methods, and classification parameters, we selected the best-performing classifier comprising 100 upregulated and 100 downregulated genes in latent parasites (**Supplementary Table 24**).

We first evaluated classifier sensitivity using our two independent post-DHA datasets (**Table 5**). Sensitivity and specificity results demonstrated that the classifier closely recapitulated the defined latent state, distinguishing them from stage-matched normal parasites with low background. We next applied the classifier to the corresponding pre-DHA populations and identified a low frequency of latent parasites before drug exposure (<1% of early trophozoites). When we compared pre- and post-DHA latency rates, drug treatment and enrichment steps yielded a ∼50-fold increase in latent parasites among early trophozoites (**Table 5**). We detected variation between NM and LH conditions but due to the potential for bias in the isolation steps of post-DHA sample processing, we confined our assessments to directly comparing pre- and post-DHA samples.

**Table 5.**
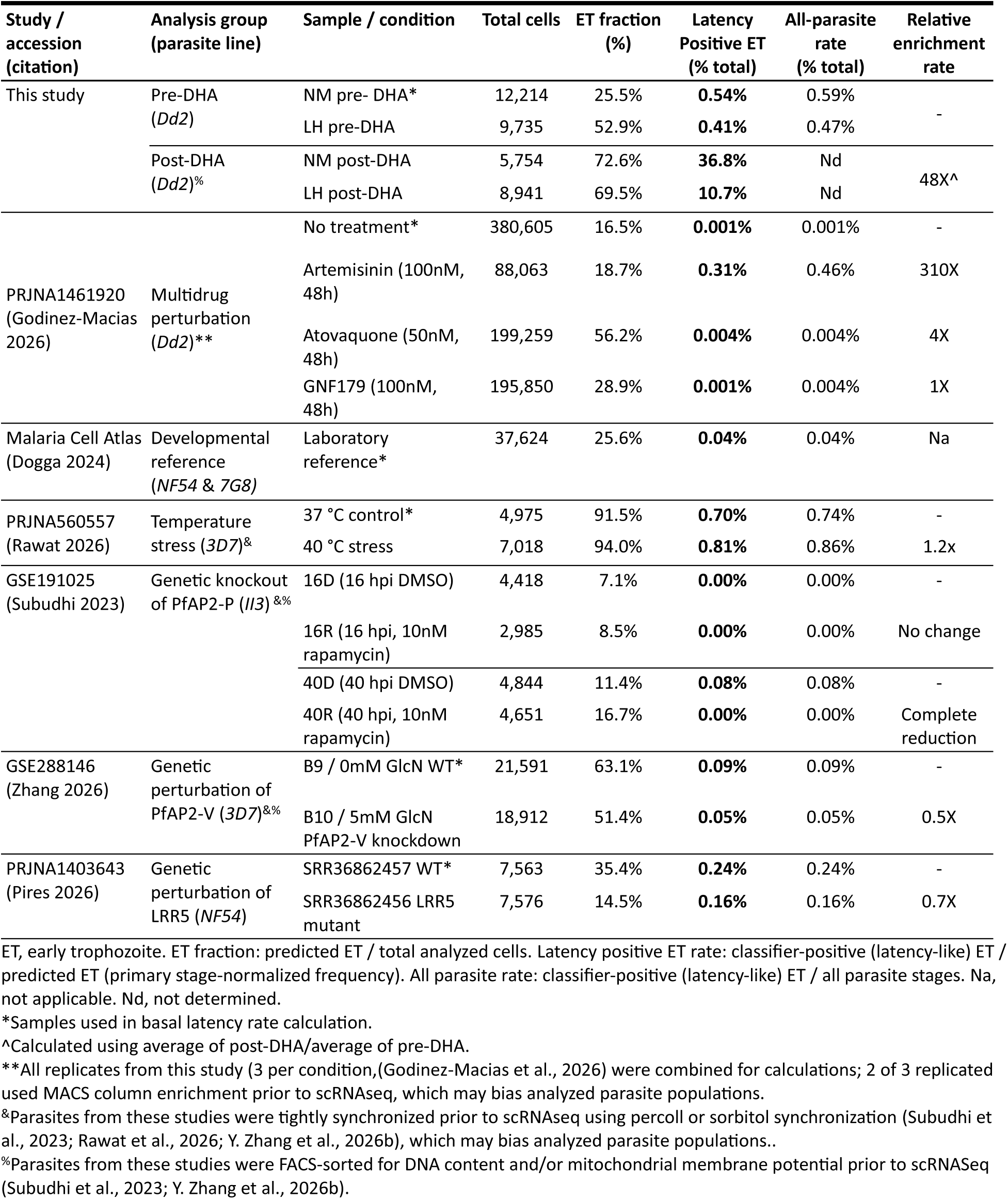
Stage-restricted application of the 200-gene latency classifier across datasets.

We also applied the classifier to a number of previously published scRNAseq datasets to assess the presence of latent-like parasites in normal and treated populations, as well as determine a basal latency rate. When assessing these data, it is important to note that we repeatedly enriched our post-DHA samples to elevate the proportion of latent compared to other stage-matched parasites (**Fig. 1A**, >10% of early trophozoite populations, **Table 5**); however, this step was not performed in other studies so latency rates are expected to be low. In one dataset comprising control, artemisinin-, and atovaquone-treated parasites (Godinez-Macias et al., 2026), we identified a strong induction of latency uniquely in the artemisinin treated condition (>300-fold). We also applied the classifier to additional independent single-cell datasets including a broad developmental reference, the Malaria Cell Atlas (Dogga et al., 2024), temperature stress conditions (Rawat et al., 2026), and several genetic perturbation datasets (Subudhi et al., 2023; Pires et al., 2026; Y. Zhang et al., 2026a). Across these datasets, the classifier identified few latent signatures, with latency rates well below 1% in most populations (**Table 5**). These findings indicate a low background frequency of transcriptionally latency-like cells across diverse parasite populations. We calculated the basal latency rate across all untreated/wild-type parasite lines at ∼0.2% of all parasites, which is relatively consistent with the level detected in our pre-DHA data set (∼0.6%). This was also in line with previously reported rates after drug exposure (range between 0.044% to 1.313%) (Teuscher et al., 2010a; Witkowski et al., 2010). However, variation in this number across diverse datasets may be due to distinct parasite lines, media conditions, and health of parasites prior to scRNAseq.

## Discussion

In this study, we show that nutrient deprivation increases the frequency of parasite entry into latency following DHA exposure (**Fig. 1A & B**) and define the transcriptional features of this state (**Fig. 5**). By integrating latency enrichment with single-cell transcriptomic analyses, we demonstrated that latency represents a distinct, actively maintained state (**Fig. 4**) that deviates from a standard developmental path. Latent parasites displayed a nearly balanced numbers of gene upregulation and downregulation (**Supplementary Tables 8 & 9**), arguing against a global shutdown. Our direct comparison of latent to stage-matched parasites revealed differences that would likely remain obscured in populations of mixed developmental stage. We found that latent parasites selectively remodeled specific biological processes involved in stress sensing, metabolic homeostasis, redox regulation, and host-cell remodeling (**Figs. 5-7**). A comparatively small effect of nutrient deprivation on the transcriptional profile of latent parasites (**Fig. 8**) suggested that environmental stress primarily influences latency entry but not the core features. A classifier built from the latent transcriptome applied to independent single-cell datasets showed that the latency program is not unique to our experimental system and enriched by artemisinin treatment in other contexts (**Table 5**). Our study supports a model in which environmental stress promotes entry into a conserved latent state characterized by preferential preservation of functions associated with survival, maintenance, and eventual reactivation.

### Environmental stress promotes entry into a distinct latent state

Previous studies of artemisinin survival have often interpreted surviving parasites primarily through the lens of dormancy, developmental delay, or proliferative arrest (Teuscher et al., 2010b; Peatey et al., 2015; Tripathi et al., 2024). Our findings refine this view by showing that latent parasites occupy a transcriptionally distinct state; reference-based mapping identified a divergent early trophozoite population that emerged after DHA exposure (**Fig. 4E & F**), and latent parasites displayed extensive transcriptional remodeling despite remaining developmentally aligned with early trophozoites. Notably, we identified a small number of latent-like parasites in pre-DHA datasets (**Table 5**), indicating that features of the latent transcriptional program may exist within a rare subpopulation prior to drug exposure. This observation raises the possibility that DHA does not exclusively induce latency itself, but instead enriches parasites already occupying, or predisposed to enter, this state. While our data cannot definitively distinguish between the two possibilities, the increase of parasites with the latency transcriptional signature following DHA exposure in independent studies demonstrates that survival is associated with a coordinated program rather than a passive consequence of growth arrest. Additionally, the presence of latent-like parasites before treatment may represent a pre-existing survival strategy that becomes advantageous under drug pressure.

Although nutrient stress is known to induce cell-cycle slowing and arrest under amino-acid limitation (Babbitt et al., 2012), we did not observe slowed development or direct entry into latency under hypoxanthine deprivation alone (**Supplementary Table 2**) (Brown et al., 2023). Instead, nutrient deprivation acted as a priming signal that increased the probability of entering latency after drug treatment (**Fig. 2A & 2B**). This increase in latency frequency was accompanied by increased long-term survival following DHA exposure, consistent with our previous nutrient-priming studies (Brown et al., 2023). However, once latency was established, latent parasites from NM and LH conditions exhibited remarkably similar transcriptional programs (**Fig. 8**), indicating that nutrient stress primarily influences entry into latency rather than maintenance of the latent state itself. These observations suggest that environmental stress influences how frequently parasites enter latency, while exerting comparatively little effect on the core transcriptional features of the state.

### Latency retains stress sensing despite limited eIF2α activation

*P. falciparum* lacks several canonical nutrient-sensing pathways found in higher eukaryotes (McLean & Jacobs-Lorena, 2017), including the mTOR signaling network that coordinates growth with nutrient availability (Ward et al., 2004). Consequently, the mechanisms by which environmental stress influences parasite survival remain incompletely understood. Our observation that nutrient limitation increases latency frequency raises the question of how environmental signals are integrated to influence survival decisions in *P. falciparum*. Despite evidence that translational repression contributes to stress adaptation and artemisinin survival in malaria parasites (Babbitt et al., 2012; Marreiros et al., 2023; M. Zhang et al., 2017), we show that enhanced latency formation occurs in the absence of eIF2α pathway activation (**Fig. 2**). However, a lack of bulk eIF2α phosphorylation following nutrient deprivation does not exclude a role for translational control in latency. eIK1-mediated eIF2α phosphorylation may occur transiently, within a restricted parasite subpopulation, or at levels below the sensitivity of population-level measurements. Indeed, our transcriptomic analyses revealed continued engagement of components associated with translational regulation (**Table 2**), alongside amino acid sensing, proteostasis, and autophagy-related processes (**Table 2**, **Figs. 5-7**). Notably, latent parasites increased expression of key eIF2α regulators, including eIK1 (Marreiros et al., 2023), together with signaling components such as PI3K and VPS15 (Bourgeois et al., 2026), suggesting coordinated engagement of multiple adaptive stress-response pathways. Rather than pointing to a single master regulator, our data support a model in which multiple stress-responsive networks converge to promote parasite persistence following drug exposure.

### Latency prioritizes cellular maintenance over growth-associated functions

The transcriptional architecture of latent parasites is consistent with a redistribution of resources away from active growth and toward cellular maintenance, stress adaptation, and long-term survival (**Figs. 5-7**, **Table 2**). The shift away from growth-associated functions extended to RNA metabolism and translational machinery; latent parasites broadly reduced expression of genes involved in RNA binding, translation initiation, ribosome biogenesis, and post-transcriptional regulation (**Fig. 5B-E**), which resembled the translationally repressed states reported during amino acid starvation (Babbitt et al., 2012; McLean & Jacobs-Lorena, 2017) and DHA-induction of dormancy (Tripathi et al., 2024). This recurrence of translational suppression across studies suggests that reduced investment in protein synthesis may represent a common feature of parasite survival rather than a response unique to any individual stressor. This idea is further supported by our comparisons between NM and LH conditions (**Fig. 8**), where reduced expression of translational machinery was largely conserved across latent parasites despite differing environmental histories.

Our comparison of latent and staged-matched parasites across multiple media conditions further revealed remodeling of downstream cellular pathways. Proteostasis pathways exhibited a pattern similar to that observed for translational machinery. Latent parasites strongly reduced transcripts to support folding newly synthesized proteins through the TRiC/CCT chaperonin complex, prefoldin-associated proteins, and multiple components of ER protein-folding machinery (**Fig. 5B-E**, **Table 2**). However, this change was not accompanied by complete loss of protein quality-control capacity. Instead, core ER, ERAD, and PROSC-associated networks exhibited mixed transcriptional regulation, with preservation of selected degradation and quality-control components despite broad reductions elsewhere in the proteostasis network (**Table 2**). This observation is partially consistent with previous studies linking artemisinin survival to enhanced proteostasis capacity and increased dependence on protein quality-control pathways (Dogovski et al., 2015; Rocamora et al., 2018). Rather than a simple increase or decrease in proteostasis activity, latency may selectively retain transcription of protein surveillance and homeostasis components.

Autophagy-associated pathways displayed pathway remodeling where latent parasites oppositely regulated transcripts encoding the autophagy-initiation partners ATG5 and ATG12 (Pang et al., 2019) (**Table 2**). Selective regulation of autophagy during latency is consistent with our previous findings that ATG8 does not mediate the enhanced survival phenotype observed following nutrient deprivation (Brown et al., 2023), while remaining partially consistent with reports linking autophagy-associated pathways to artemisinin survival (Ray et al., 2022b). Our findings may help reconcile these observations; latency is accompanied by engagement of some autophagy-related functions rather than uniform activation of the pathway. Similar non-uniform regulation of autophagy has been reported in diverse biological systems, where specific functions can be maintained rather than wholesale activation of the pathway (Lamark & Johansen, 2021).

We also observed a selective remodeling of mitochondrial and redox metabolism. Despite broader reductions in metabolism-associated transcripts (**Fig. 6**), latent parasites retained key transcripts associated with mitochondrial maintenance and cellular homeostasis. In particular, retention of electron transport chain (ETC)-associated functions alongside remodeling of specific metabolic pathways suggested continued investment in processes required for long-term viability. This pattern aligns with earlier observations that recovery from DHA-induced dormancy depends on maintenance of mitochondrial function (Peatey et al., 2015). It is also consistent with recent transcriptomic analyses showing that dormant parasites preserved selected metabolic activities despite extensive suppression of growth-associated processes (Tripathi et al., 2024). Likewise, latent parasites largely preserved components supporting glutathione production and NADPH generation despite reductions in glutathione turnover pathways, indicating continued investment in antioxidant capacity and redox homeostasis during latency. Coupled with the retention of ETC-associated activities, these observations suggest that latent parasites maintain metabolic pathways required to sustain cellular integrity during prolonged periods of growth suppression and facilitate eventual reactivation. Such preservation may be particularly important in the context of artemisinin-induced oxidative and proteotoxic stress, which promotes protein damage and engagement of cellular quality-control pathways (Dogovski et al., 2015; Rocamora et al., 2018).

One of the clearest examples of pathway-level uncoupling during the latent state involved hemoglobin metabolism. Latent parasites maintained expression of components associated with hemoglobin uptake, including K13 and multiple KIC proteins, while reducing expression of proteases involved in downstream hemoglobin digestion (**Fig. 6**). To our knowledge, this dissociation between hemoglobin uptake and digestion has not been previously associated with parasite latency. Reduced hemoglobin digestion has been repeatedly implicated in artemisinin resistance through decreased drug activation (Klonis et al., 2013; Mok et al., 2015; Bhattacharjee et al., 2018; Yang et al., 2019). The preservation of uptake alongside reduced digestive capacity during latency could simultaneously restrict heme release and artemisinin activation while preserving access to resources for eventual reactivation.

Finally, preservation was also evident across broader metabolic networks. In addition to the maintenance of selected mitochondrial, redox-associated, and signaling functions discussed above, latent parasites exhibited coordinated upregulation of multiple acyl-CoA synthetases (**Fig. 6**). In fact, several of the ACS family members were among the most strongly induced genes in the latent transcriptome. This coordinated induction of multiple ACS family members is notable because previous studies have shown that each member of the expanded ACS gene family likely performs specialization functions (Matesanz et al., 2003). Although the functions of ACS1-9 remain incompletely resolved, acyl-CoA synthetases generally activate fatty acids for downstream use. This function, combined with a strong upregulation of the lipid scavenging protein XLH4 (Liu et al., 2024), points to a previously unrecognized role for fatty-acid activation and lipid handling during latency. Additionally, this observation is particularly intriguing given emerging evidence that ACS-family enzymes, including ACS10 and ACS11, influence antimalarial drug susceptibility and parasite lipid metabolism (Bopp et al., 2023). Together, these findings raise the possibility that lipid utilization is among the limited set of functions preserved during latency.

### Latency preserves host-cell remodeling and erythrocyte maintenance

A particularly unexpected feature of latent parasites was the continued investment in host-cell remodeling despite broad suppression of growth-associated programs (**Table 3**, **Fig. 7**). While latent parasites downregulated *PfEMP1* expression when compared to their stage-matched non-latent counterparts (**Fig. 7B**), they maintained or upregulated numerous components of protein export and erythrocyte remodeling machinery. These included PTPs, other Maurer’s cleft-associated proteins, PHIST family members, and HSP40 co-chaperones (**Fig. 7D**). Because PTPs are critical for *PfEMP1* expression on the erythrocyte surface, proper knob formation, and extracellular vesicle release from the Maurer’s cleft (Carmo et al., 2022), their retention despite reduced *PfEMP1* expression suggests that latent parasites selectively preserved aspects of the export machinery while altering its functional output.

We also observed additional evidence of maintained host-cell investment by latent parasites through the upregulation of RESA and FIKK family transcripts (**Fig. 7C & 7D**). RESA proteins maintain erythrocyte integrity from heat-shock induced damage (Da Silva et al., 1994; Diez Silva et al., 2005) while both FIKK kinases and RESA proteins reorganize the erythrocyte cytoskeleton to address membrane rigidity and knob formation (Mills et al., 2007; Diez-Silva et al., 2012; Zhang et al., 2015). Because erythrocyte rigidity and cytoskeleton organization influence parasite circulation and splenic clearance (Suwanarusk et al. 2004), maintenance of these remodeling proteins may help preserve a suitable host-cell environment during prolonged periods of growth suppression.

Interestingly, latent parasites upregulate other antigen families including SURFs, SERAs, and MSPs, suggesting that export machinery may support their transport to the host erythrocyte surface. Although functional studies will be required to determine whether these changes alter immune evasion, sequestration, or rosetting behavior, the overall transcriptional pattern suggests that latent parasites continue investing in maintenance of the infected erythrocyte rather than abandoning host-cell remodeling altogether. To our knowledge, erythrocyte maintenance by latent parasites has not previously been described in *P. falciparum* and represents another example of selective preservation during latency.

### Latency is a reversible state poised for reactivation

An important feature of malaria parasite biology is that parasites recovering from artemisinin treatment remain susceptible to subsequent drug exposure (Ittarat et al., 2003). This supports the existence of a rapid, reversible adaptive response that promotes survival of acute drug stress without commitment to a permanently resistant state. The recovery kinetics observed here further support this view, suggesting that latency represents a transient physiological state rather than a terminal endpoint. By day three post-treatment (**Fig. 1**), we observe in vitro populations consistent with both latency establishment and reactivation (**Fig. 4**). The coexistence of these populations indicates that latency is not a terminal endpoint but a dynamic transitional state through which parasites pass as they recover from drug exposure. Similar strategies have been described in other *Plasmodium* quiescent states, including *P. cynomolgi* hypnozoites (Voorberg-van der Wel et al., 2017) and nutritionally stressed mosquito-stage oocysts (Habtewold et al., 2021), where prolonged reductions in growth are accompanied by preservation of future developmental potential. Several features of the latent transcriptional program are consistent with such a poised state. Despite broad reductions in RNA-processing and translation-associated pathways, latent parasites retained expression of invasion-associated genes, including multiple MSP and EBA family members, together with mitochondrial maintenance functions and nutrient acquisition pathways. The preservation of these functions is consistent with a state that remains prepared for future growth and host-cell invasion despite temporary suppression of proliferation.

### Latency is distinct from canonical artemisinin resistance

Our previous work demonstrated that nutrient-priming-induced drug tolerance occurs across multiple parasite genetic backgrounds, including parasites differing in K13 mutation status (Brown et al., 2023). The detection of latency-associated transcriptional signatures across independent datasets further suggests that this state is not restricted to our experimental system (**Table 5**). Latent-like parasites were detected in both untreated populations and parasites exposed to a range of stress conditions, suggesting that latency represents a general parasite stress-response program rather than a response unique to DHA exposure. The presence of latent-like parasites before drug treatment also raises the possibility that latency contributes to persistence by allowing a subset of parasites to survive transient environmental stress without requiring stable resistance mutations. However, the relationship between latency and K13-mediated artemisinin resistance remains unresolved. K13 mutations are thought to promote survival, in part, through altered endocytosis and reduced hemoglobin catabolism, thereby limiting heme-dependent activation of DHA (Birnbaum et al., 2020). In contrast, latent parasites exhibited increased expression of K13 and multiple KIC-associated uptake components while simultaneously reducing downstream digestive functions (**Fig. 6**). This pattern suggests that latency and K13-mediated resistance may represent different strategies for surviving artemisinin exposure. Whether K13 genotype influences the frequency of latency formation, maintenance, or reactivation remains unknown and will require direct comparison of K13 wild-type and mutant parasites under matched drug exposure conditions.

### Challenges and opportunities for defining malaria latency

A major strength of this study is the direct analysis of samples enriched in latent parasites. By removing actively growing parasites prior to scRNAseq analysis, we minimized transcriptional signals arising from parasites that had already resumed development or survived DHA exposure without entering latency. Profiling parasites three days after DHA treatment captured a critical transitional window in which latency remained established while a subset of parasites had already begun to reactivate. This experimental design enabled direct comparison of latent and stage-matched non-latent parasites, revealing coordinated pathway remodeling that would likely remain obscured in mixed populations or analyses that do not account for developmental stage.

Despite this strength, some limitations should be considered when interpreting our findings. First, many conclusions are derived from transcriptional profiles and therefore do not establish functional activity of the underlying pathways. Although the observed patterns are highly consistent across multiple analyses, experimental validation will be required to determine whether the inferred pathway changes translate into changes at the protein level.

Second, the current study captures latency at a single post-treatment window and therefore cannot fully resolve the molecular programs of entry, maintenance, or reactivation. While comparisons between NM- and LH-derived latent parasites suggest that environmental stress primarily influences latency entry rather than maintenance (**Fig. 8**), definitive separation of these stages will require longitudinal profiling across the full trajectory from drug exposure through recovery. Such experiments could help distinguish pathways that initiate latency from those that sustain or terminate the latent state.

Third, our observations differ in some respects from studies that identified early ring stages as the primary artemisinin-tolerant population (Auparakkitanon & Wilairat, 2023; Platon et al., 2023). These studies largely rely on ring-stage survival assays that examine parasite survival during the first hours following invasion, whereas our analyses focused on parasites several days after DHA exposure. This difference in experimental timing likely contributes to the identification of a transcriptionally active latent population occupying early trophozoite transcriptional space rather than a ring-stage-restricted state. Although our enrichment strategy may impact recovery of early ring-stage parasites, particularly those that fail to maintain viability, the localization of latent cells within trophozoite transcriptional space (**Fig. 4**), together with independent stage marker validation (**Supplementary Fig. 5C**), supports the conclusion that the population characterized in the current study represents early trophozoites rather than misclassified ring stages.

Finally, some metabolic features observed in latent parasites partially overlap with transcriptional responses previously reported under low-hypoxanthine conditions (Tewari et al., 2019). While the strong conservation of latent transcriptional programs between NM and LH conditions argues against nutrient limitation as the primary driver of the latent phenotype (**Fig. 8**), we cannot exclude the possibility that certain metabolic signatures reflect contributions from prior nutrient history. Future studies comparing multiple environmental stressors and nutrient perturbations will help distinguish core features of latency from responses unique to individual priming conditions.

### Future directions for understanding malaria latency

Interventions that disrupt latency establishment, maintenance, or reactivation may complement existing antimalarial therapies that primarily target actively proliferating parasites. Several observations from this study identify promising avenues for future investigation. The retetion of TCA cycle, redox homeostasis, stress signaling, autophagy, and host cell remodeling pathways suggests that these functions may contribute to latency establishment or maintenance. Similarly, continued expression of invasion-associated genes and erythrocyte-remodeling factors raises the possibility that latent parasites actively prepare for future proliferation while modifying the host-cell environment to support long-term survival. Determining whether these transcriptional programs influence parasite fitness, host-cell remodeling, immune recognition, or reactivation kinetics will be important next steps.

More broadly, this work demonstrates the value of combining physiological perturbation with single-cell transcriptomics to uncover adaptive survival states that remain obscured in bulk population measurements. By revealing latency as a poised and transcriptionally organized survival strategy, these findings provide a framework for understanding how malaria parasites persist following drug exposure and identify new opportunities to target survival mechanisms that operate outside of conventional resistance pathways.

## Methods

### Parasite culture and nutrient-priming conditions

We obtained *Plasmodium falciparum* line MRA-156 (*Dd2*) from the Malaria Research and Reference Resource Center (MR4 BEI Resources). We cultured the parasites in A+ human erythrocytes (Valley Biomedical, Winchester, VA) at 3% hematocrit using RPMI 1640 medium buffered with HEPES (Sigma-Aldrich, St. Louis, MO), supplemented with 0.5% Albumax II lipid-rich BSA (Sigma-Aldrich, St. Louis, MO) and 50 mg/L hypoxanthine (Sigma-Aldrich, St. Louis, MO). We refer to this formulation as complete medium, which served as the normal-medium (NM) control condition throughout the study unless otherwise indicated.

We maintained the cultures at 37 °C and flushed them individually with a gas mixture of 5% oxygen, 5% carbon dioxide, and 90% nitrogen. We diluted cultures with uninfected erythrocytes and replaced the medium every other day. We monitored parasitemia by flow cytometry (BD Accuri C6, Ann Harbor, MI) using SYBR Green I (Thermo Fisher Scientific, Waltham, MA) and maintained it below 2% during routine culture.

To investigate how environmental stress influences latency formation, we exposed synchronized parasites to previously established mild nutrient limiting conditions that reduce growth while preserving viability (Brown et al., 2023). We used two priming conditions: low-hypoxanthine (LH) medium and thiamine-free (TF) medium. We prepared LH medium using the RPMI 1640 HEPES base medium (Sigma-Aldrich, St. Louis, MO) supplemented with reduced exogenous hypoxanthine (Sigma-Aldrich, St. Louis, MO). We diluted 2mM hypoxanthine stock solution in DMSO (Thermo Fisher Scientific, Waltham, MA) 1:4,000 to achieve the final concentration of 0.5 μM. We obtained thiamine-free (TF) RPMI medium formulated to match RPMI 1640 HEPES (Sigma-Aldrich, St. Louis, MO) without thiamine hydrochloride. Unless otherwise specified, we supplemented all low-nutrient media with 0.5% Albumax II lipid-rich BSA (Sigma-Aldrich, St. Louis, MO).

We preincubated uninfected erythrocytes in the appropriate low-nutrient medium for 48 h at 37 °C, 3% hematocrit, under a gas mixture of 5% oxygen, 5% carbon dioxide, and 90% nitrogen (Brown et al., 2023). We handled corresponding NM control erythrocytes identically. After preincubation in the corresponding medium, we initiated priming experiments by combining uninfected erythrocytes with infected cultures and adjusting starting parasitemia to 0.1% for LH experiments and 0.1-0.25% for TF experiments. For LH condition experiments, we refreshed the culture medium at 48h and for TF, at 72h (Brown et al., 2023). We collected aliquots for flow cytometry using flow cytometry using SYBR Green I and MitoProbe DiIC1(5) as described below (see *Enrichment and characterization of latent parasites*). For nutrient deprivation experiments, we maintained parasite cultures at 37 °C under the same gas conditions, diluted cultures with uninfected preincubated erythrocytes, and replaced the culture medium every other day.

We compared nutrient-deprived samples with NM to assess growth, mitochondrial membrane potential (MMP) status, and ring-stage proportions. Nutrient deprivation was mild due to previously defined levels of growth reduction and minimal impact on mitochondrial membrane potential (Brown et al., 2023). We considered LH nutrient deprivation successful when cultures met the following criteria relative to controls: (i) approximately >10% reduction in growth, (ii) minimal reduction in MMP (∼10% decrease), and (iii) ring-stage percentages within ∼15% of non-primed cultures. For TF conditions, the criteria for success included only (i) and (ii). We accepted a broad range of ring-stage proportions relative to NM controls (<15%).

### Generation of latent parasites following DHA exposure

We synchronized parasites using 5% D-sorbitol (Thermo Fisher Scientific, Waltham, MA) following the method described previously (Lambros & Vanderberg, 1979b). Following synchronization, we exposed parasites to NM, LH, or TF conditions (see *Parasite culture and nutrient-priming conditions*) before treating cultures with DHA. To distinguish effects on parasite recovery from latency formation, we treated cultures with either 200nM dihydroartemisinin (DHA; Selleck Chemicals, Houston, TX) (growth-recovery experiments (Brown et al., 2023) or 700nM DHA (latency-enrichment experiments, (Witkowski et al., 2013). Following 6h of DHA exposure, we removed drug by washing cultures three times in complete medium before returning cultures to standard growth conditions. We assessed parasite staging by flow cytometry using SYBR Green I and MitoProbe DiIC1(5) as described below (see *Enrichment and characterization of latent parasites*).

### Enrichment and characterization of latent parasites

To enrich latent parasites prior to scRNAseq, we depleted actively growing parasites by magnetic column separation (MACS) as described previously (Ribaut et al., 2008), using LS columns, a magnetic separator (Miltenyi Biotec, Gaithersburg, MA), and a three-way-stopcock (Smiths Medical, Dublin, OH) to control the flow speed. Columns retained mature parasite-infected erythrocytes containing hemozoin, whereas uninfected erythrocytes, early-stage parasites, and nonviable cells passed through. By collecting the flow-through fraction, we enriched the population for latent parasites while reducing contributions from actively recovering parasitesto the downstream scRNAseq analyses. During MACS, we diluted the infected erythrocyte cultures 1:2 in complete medium prior to two rounds of column loading to reduce hematocrit and facilitate column flow. Following binding, we washed the column with prewarmed medium to remove residual unbound cells and pooled the flow-through fractions before returning to the incubator for staining.

We used flow cytometry to monitor parasitemia, mitochondrial membrane potential, and parasite viability and to isolate live parasites for single-cell sequencing. We stained parasites using SYBR Green I (Thermo Fisher Scientific, Waltham, MA, 1:10,000 in 1X PBS) and MitoProbe DiIC₁(5) (Thermo Fisher Scientific, Waltham, MA, 1:400 in 1XPBS) under a gas mixture of 5% oxygen, 5% carbon dioxide, and 90% nitrogen before dilution in 1XPBS and analysis by flow cytometry (50,000 events per sample unless otherwise indicated).

### Flow sorting, single-cell library preparation, and sequencing

For parasite isolation, we sorted SYBR Green I-positive and MitoProbe DiIC₁(5)-positive parasites using a Sony SH800S cell sorter (Sony Biotechnology, San Jose, CA). We conducted parasite staining as for flow cytometry except that we resuspended parasites in RPMI 1640 HEPES supplemented with 1% bovine serum albumin (BSA, Thermo Fisher Scientific, Waltham, MA). For sorting, we used 70μm microfluidic chips (Sony Biotechnology, San Jose, CA), and gated and sorted viable parasites, collecting 40,000 parasites per treatment condition.

Because *Plasmodium*-infected erythrocytes contain low RNA content and are sensitive to extended processing, we optimized sorting conditions by comparing parasitemia and mitochondrial membrane potential before sorting and at 1h and 6h after sorting (**Supplementary Table 3**).

For scRNAseq, we loaded sorted parasite-containing erythrocytes onto the 10x Genomics Chromium platform using the Chromium Next GEM Single Cell 3′ Gene Expression v3.1 workflow and Chromium Chip G according to the manufacturer’s instructions. We generated libraries using the standard 10x 3′ gene-expression workflow, including single-cell partitioning, gel bead-based barcoding, reverse transcription, cDNA amplification, fragmentation, adapter ligation, sample index PCR, and library cleanup. We assessed library size and quality using an Agilent TapeStation and pooled indexed libraries and sequenced on an Illumina platform using paired-end sequencing and the standard 10x 3′ read structure. We then targeted sequencing depth to recover sufficient parasite transcript complexity for downstream clustering, reference mapping, and differential-expression analyses.

### Single-cell preprocessing, integration, and developmental stage annotation

We converted raw sequencing data to FASTQ files and processed them using Cell Ranger Count. We demultiplexed and aligned reads, assigned them to genes, and collapsed information into gene-by-cell unique molecular identifier (UMI) count matrices using the Cell Ranger gene-expression pipeline. We generated a *P. falciparum* 3D7 reference from PlasmoDB release 68 genome and annotation files and used it for alignment and gene quantification. We generated Cell Ranger outputs independently for each sample including downstream sample-level integration and comparative analyses. We imported filtered feature-barcode matrices from Cell Ranger into R and generated Seurat objects for each sample. We removed genes detected in very few cells and excluded cells with insufficient genes or UMIs before downstream analysis. We examined parasite mitochondrial transcript content and library complexity metrics as quality-control features and retained cells passing quality-control thresholds for further analysis. We used Seurat to perform normalization, dimensionality reduction, clustering, and integration. We analyzed pre-DHA and post-DHA datasets both individually and as combined integrated objects to distinguish condition-specific effects from shared developmental and latency-associated structure. We used integrated UMAP embeddings to compare NM and LH conditions, assess overlap between pre- and post-DHA populations, and identify drug-associated transcriptional branches.

We projected parasite transcriptomes onto the *P. falciparum* Malaria Cell Atlas reference (Howick et al., 2019) to assign developmental-stage identities including early and late ring, trophozoite, schizont, sexual-commitment, and gametocyte-associated states. We validated stage assignments by examining curated marker genes for ring, trophozoite, schizont, and sexual-stage populations (see *Identification and validation of latent parasites*).

### Identification, validation, and characterization of latent parasites

We identified latent parasites from post-DHA datasets by locating a DHA-specific branch that diverged from the normal early trophozoite trajectory. This branch was absent from matched pre-DHA samples, so we designated it as the latent population. We validated latent-population assignments using three complementary approaches: (i) sexual-stage contamination analysis, (ii) reference-annotation confidence scores, and (iii) data-driven stage-signature projection analyses.

We evaluated the potential for sexual-stage contamination using curated sexual-commitment and gametocyte marker scores, using a threshold based on the upper tail of the normal early trophozoite sexual-marker distribution was used to identify sexual-marker-high cells. We examined prediction-score distributions from the reference annotation model to determine whether latent cells retained high-confidence early trophozoite identity. We generated data-driven stage signatures from normal stage populations and projected onto latent cells to test whether latent trophozoites retained the full normal early trophozoite transcriptional program or instead resembled another developmental stage. We used developmental-adjacency analyses to classify strong alternative-stage signals as self, adjacent developmental continuum, or distant contamination.

To define the biological features of latency, we compared latent and stage-matched normal early trophozoites using module scoring, differential-expression analysis, and pathway-enrichment approaches. We used curated developmental, metabolic, stress-response, and sexual-stage gene sets to quantify pathway activity and validate cell identities.

We computed module scores per cell using Seurat AddModuleScore and, in parallel, as the z-scaled mean expression of module genes. We used three classes of curated gene set: developmental-stage markers for annotation validation, a sexual-commitment/gametocyte marker set for contamination assessment, and curated metabolic modules for metabolic-state analysis. In addition to curated marker sets, we derived data-driven stage signatures directly from the dataset as described above. Gene-set membership is listed in **Supplementary Table 6**.

We scored latent and normal early trophozoites for curated metabolic gene modules using both background-adjusted AddModuleScore and z-scaled mean expression. We added an initial glycolysis/lactate module as first test of glycolytic suppression and used an expanded 52-gene metabolism module to summarize broader metabolic activity. We then compared pathway-resolved modules between latent and normal early trophozoites, including glycolysis/lactate, TCA cycle, fumarate/aspartate metabolism, cytosolic oxaloacetate-to-lactate metabolism, mitochondrial respiration, glutamine/glutamate metabolism, glutathione/redox metabolism, pentose phosphate pathway, and pyridoxal phosphate synthesis.

We calculated median module scores for latent and normal early trophozoites within each post-DHA dataset. We interpreted differences as latent minus normal, such that negative values indicated lower module activity in latent cells. We used concordant results between AddModuleScore and z-scaled mean expression to support robust pathway-level differences.

We performed differential-expression analysis to compare latent early trophozoites with stage-matched normal early trophozoites using Model-based Analysis of Single-cell Transcriptomics (MAST) (Finak et al., 2015). We combined latent cells from post-DHA NM and LH conditions for the primary latent-versus-stage-matched comparison because DHA exposure represented the shared condition. We calculated Log_2_ fold change and adjusted p-value and defined high-confidence differentially expressed genes using an adjusted p-value < 0.05 and an absolute Log_2_ fold-change ≥ 1. We interpreted upregulated genes as enriched in latent parasites, while downregulated genes were interpreted as depleted in latent parasites relative to normal trophozoites.

We performed gene set enrichment analysis (GSEA) using the fgsea package (v. 1.36.2). We obtained the 3D7 genome sequence for alignment and annotation, as well as gene product descriptions and metabolic pathway information from PlasmoDB release 68 and supplemented it with a curated artemisinin tolerance gene set (Brown et al., 2023) and an additional manually curated DNA repair gene set. We conducted visualizations and downstream analyses in R (v. 4.5.2) using tidyverse packages (v. 2.0.0).

We performed Gene Ontology (GO) enrichment analysis on upregulated and downregulated gene sets, defined by using the same high-confidence differential expression threshold (adjusted p-value < 0.05, |log2FC| ≥ 1), using g:Profiler. We imported enrichment results into R and filtered them to retain only GO terms from the Biological Process (BP), Cellular Component (CC), and Molecular Function (MF) categories. We combined results from both directions and annotated them as upregulated or downregulated to enable direct comparison.

Within each GO category, we ranked terms by adjusted p-values and selected the most significant terms for visualization. We retained broad, high-level GO terms (e.g., “biological process”) and explicitly labeled them as broad to distinguish them from more specific functional annotations. We visualized enrichment results as horizontal bar plots using −log10 adjusted p-values and displayed the number of contributing genes for each term. We colored upregulated terms red and downregulated terms blue.

To link enriched GO terms to their underlying genes, we expanded gene lists associated with each term and merged them with a gene annotation file containing gene identifiers and gene names. We generated GO-to-gene mapping tables separately for BP, CC, and MF categories and organized genes side-by-side across terms to facilitate interpretation of shared patterns. We saved all plots and tables in a structured output directory to support reproducibility and downstream analyses.

We identified uncharacterized genes from the latent-versus-normal differential-expression table by selecting genes annotated as conserved *Plasmodium* proteins of unknown function, hypothetical proteins, exported proteins of unknown function, or related unknown-function categories. We retained genes meeting the high-confidence differential-expression threshold of adjusted p-value < 0.05 and |log2FC| ≥ 1 for downstream analysis. To assign putative functions to these genes, we analyzed PlasmoFP predictions using an effective FDR threshold of ≤ 0.05. We summarized predicted Gene Ontology terms by molecular function, biological process, and cellular component to determine whether uncharacterized latency-associated genes converged on biological themes identified from the annotated latency program.

To characterize host-parasite interaction programs during latency, we assembled curated gene sets representing PfEMP1/var genes, RIFINs, PHISTs, Maurer’s cleft proteins, and PfEMP1 trafficking proteins (PTPs) using PlasmoDB release 68 and published annotations (Bekić & Kilian, 2023). Complete gene-set membership is provided in **Supplementary Table 16**.

### Latency-signature discovery and cross-dataset validation

To construct a portable latency classifier, we used transcriptionally defined latent and normal early trophozoites from the post-DHA NM and LH datasets as labeled populations. We evaluated more than 150,000 combinations of gene-panel size, scoring method, normalization procedure, and classification threshold. We assessed candidate models using repeated 20 × 5-fold cross-validation, with threshold calibration performed within the training folds, and leave-one-dataset-out validation between NM and LH. The selected classifier comprised the 100 genes upregulated and the 100 genes downregulated in latent parasites that produced the best overall discrimination. We provide the complete 200-gene panel in **Supplementary Table 24**.

For each cell, we calculated separate scores for the 100-gene latency-up and 100-gene latency-down panels using the module-scoring framework described above, standardized the two component scores, and calculated the reciprocal latency score as standardized UP score minus standardized DOWN score; higher values indicated stronger expression of the latent program together with weaker expression of the normal early-trophozoite program. We calibrated the latency-positive threshold among stage-matched normal early trophozoites in the post-DHA NM dataset. Classifier performance was then evaluated independently in the post-DHA NM and LH datasets using the transcriptionally defined latent and normal populations, with sensitivity calculated as classifier-positive latent cells divided by all defined latent cells and false-positive rate calculated as classifier-positive normal early trophozoites divided by all stage-matched normal early trophozoites.

We applied the fixed 200-gene panel and calibrated threshold, without gene reselection, to the matched pre-DHA datasets and to independent single-cell datasets, including the Malaria Cell Atlas developmental reference and multidrug-, temperature-, and genetic-perturbation studies (**Table 5**). We first assigned developmental stages in each dataset and restricted latency-positive calls to cells annotated as early trophozoites. We visualized continuous classifier scores on UMAP embeddings and used the fixed threshold to generate binary latency-positive calls. For each dataset, the stage-normalized latency-positive early-trophozoite rate was calculated as classifier-positive early trophozoites divided by all predicted early trophozoites; the all-parasite rate was calculated as classifier-positive cells divided by all analyzed parasite cells. Signals in external datasets were described as latency-like.

### Visualization and statistical analysis

We performed all analyses in R using Seurat, tidyverse, ggplot2, fgsea, and related packages. We assessed differential expression with MAST, and pathway enrichment analyses used multiple-testing-corrected significance thresholds. We exported all gene lists, pathway assignments, module scores, and statistical outputs as supplementary files to support reproducibility.

