## Supplementary Fig. for "Environmental stress promotes entry into a pre-existing latent state in *Plasmodium falciparum*"

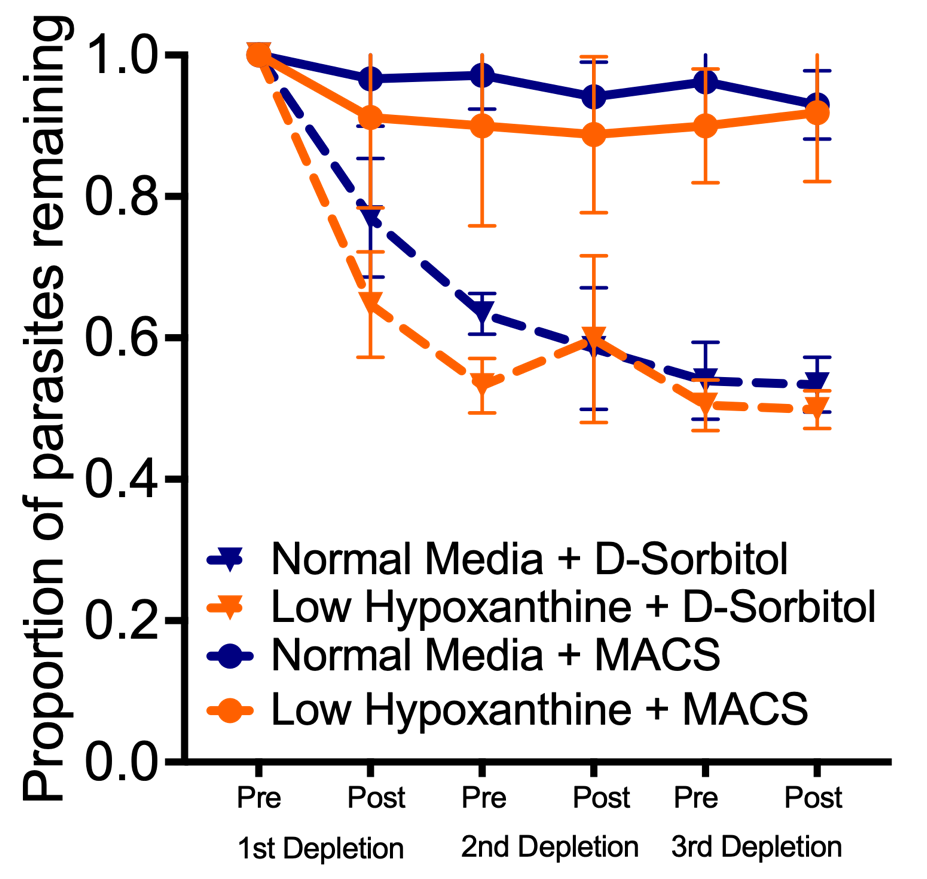

**Supplementary Figure 1. MACS preserves more parasite material than D-sorbitol lysis across successive depletions of actively growing parasites.** We depleted actively growing parasites from normal media (NM) or low hypoxanthine (LH) cultures by either D-sorbitol lysis (n = 3) or magneticly activated cell sorting (MACS, n = 6), measuring parasitemia before and after each of three depletions. We normalized parasitemia within each culture to the value measured before the first depletion (value of 1). Points show mean ± SEM.

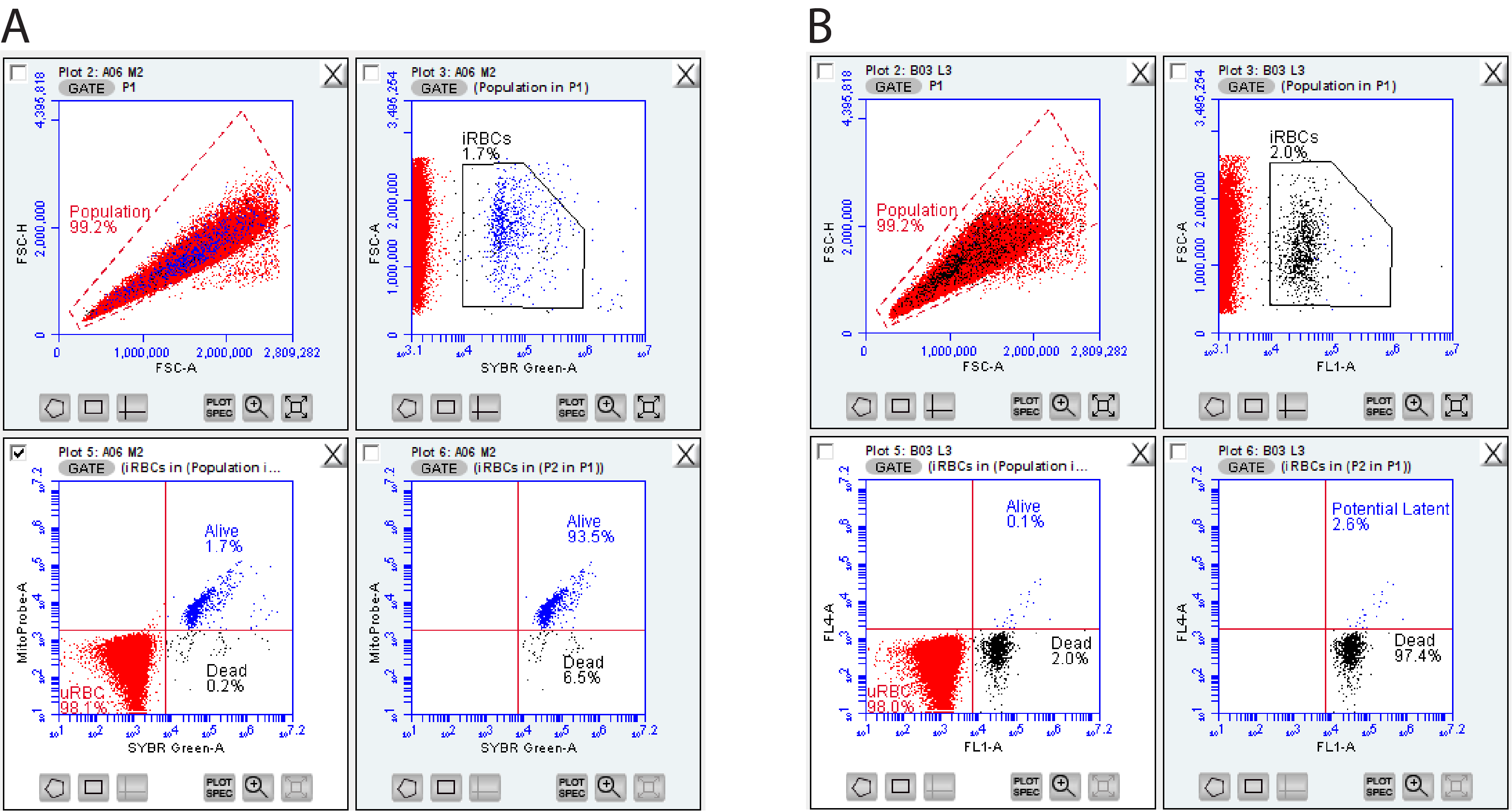
**Supplementary Figure 2. Gating strategy for flow cytometry-based quantification of parasite viability and latency.** **[A]** Representative gating for an untreated, actively growing *P. falciparum* culture. Cells are first resolved by SYBR Green into uninfected RBCs (uRBC, red) and SYBR Green-positive infected RBCs (iRBC) (Plot 3); iRBCs are further resolved by SYBR Green and MitoProbe co-staining into live (blue) and dead (black) parasites (Plot 5). Proportions of live and dead parasites are quantified from “Alive” quadrant of [A] in Plot 6, which contains only these two populations. **[B]** Representative gating showing latent parasite population from a normal medium (NM) replicate (Experiment 8, **Supplementary Table 2**), using the same quantification scheme as in [A]. Latent parasites are quantified from the "Potential Latent" quadrant of [B], Plot 6.

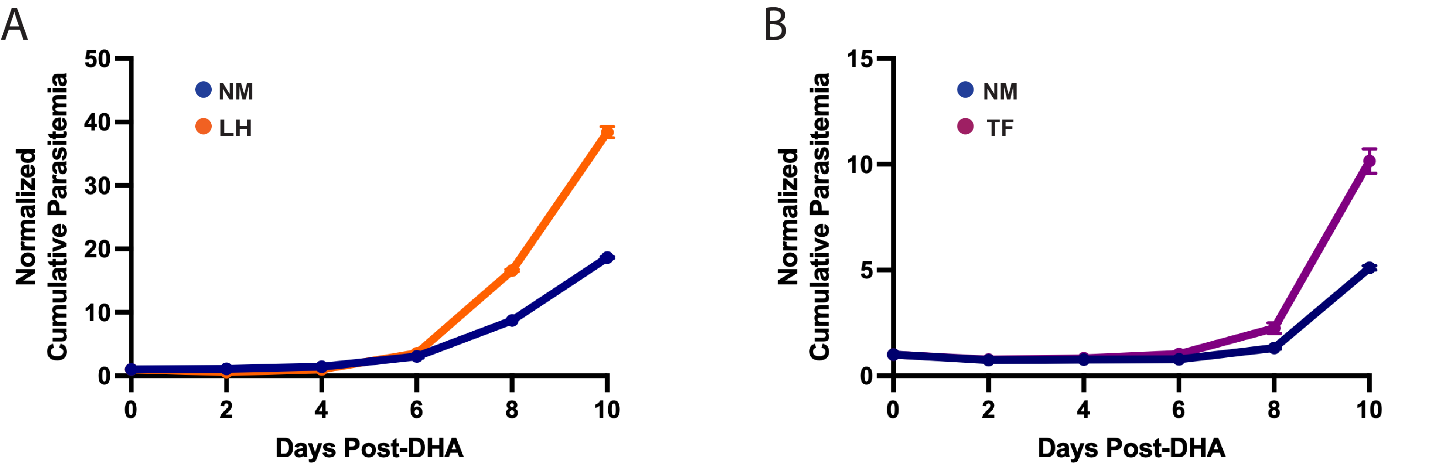
**Supplementary Figure 3. Nutrient deprivation leads to improved post-DHA recovery**. Recovery of *P. falciparum* in normal medium following deprivation in **[A]** low hypoxanthine (LH) medium or **[B]** thiamine-free (TF) medium. Error bars represent SEM of replicates from one representative experiment. Results from independent experiments are detailed in **Supplementary Table 2** (LH, n=6; TF, n=5).

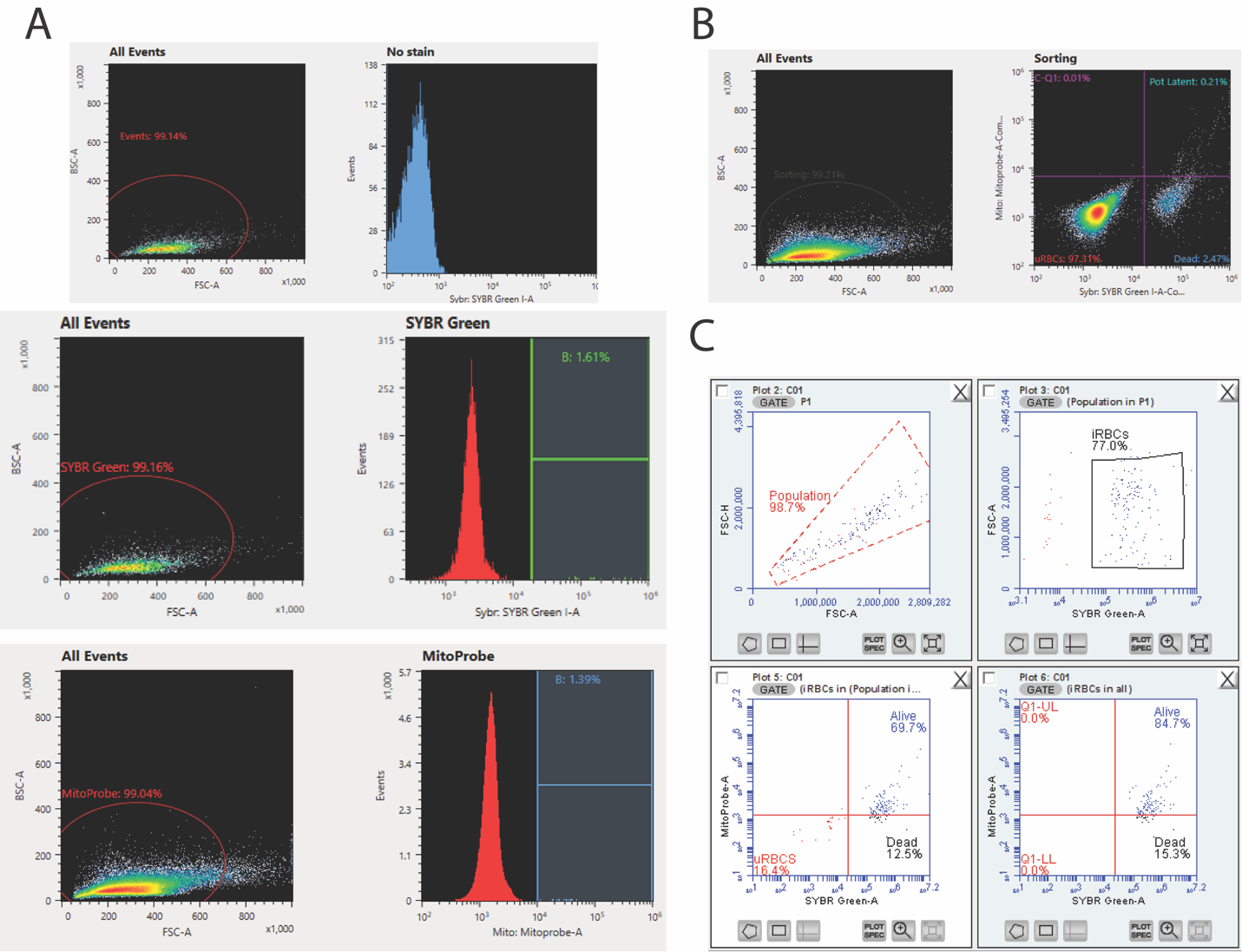

**Supplementary Figure 4. Quality control before and after FACS sorting to obtain viable latent parasites for scRNA-seq**. All plots from a normal medium (NM) as representative of FACS gating. **[A]** Compensation matrices from sample prior to sorting. **[B]** FACS gating during sorting for latent parasites using SYBR Green/MitoProbe staining after compensation. **[C]** Post-FACS quality control, using the same flow cytometry gating strategy as in Fig. S3A, confirming parasite viability following sorting in [B].

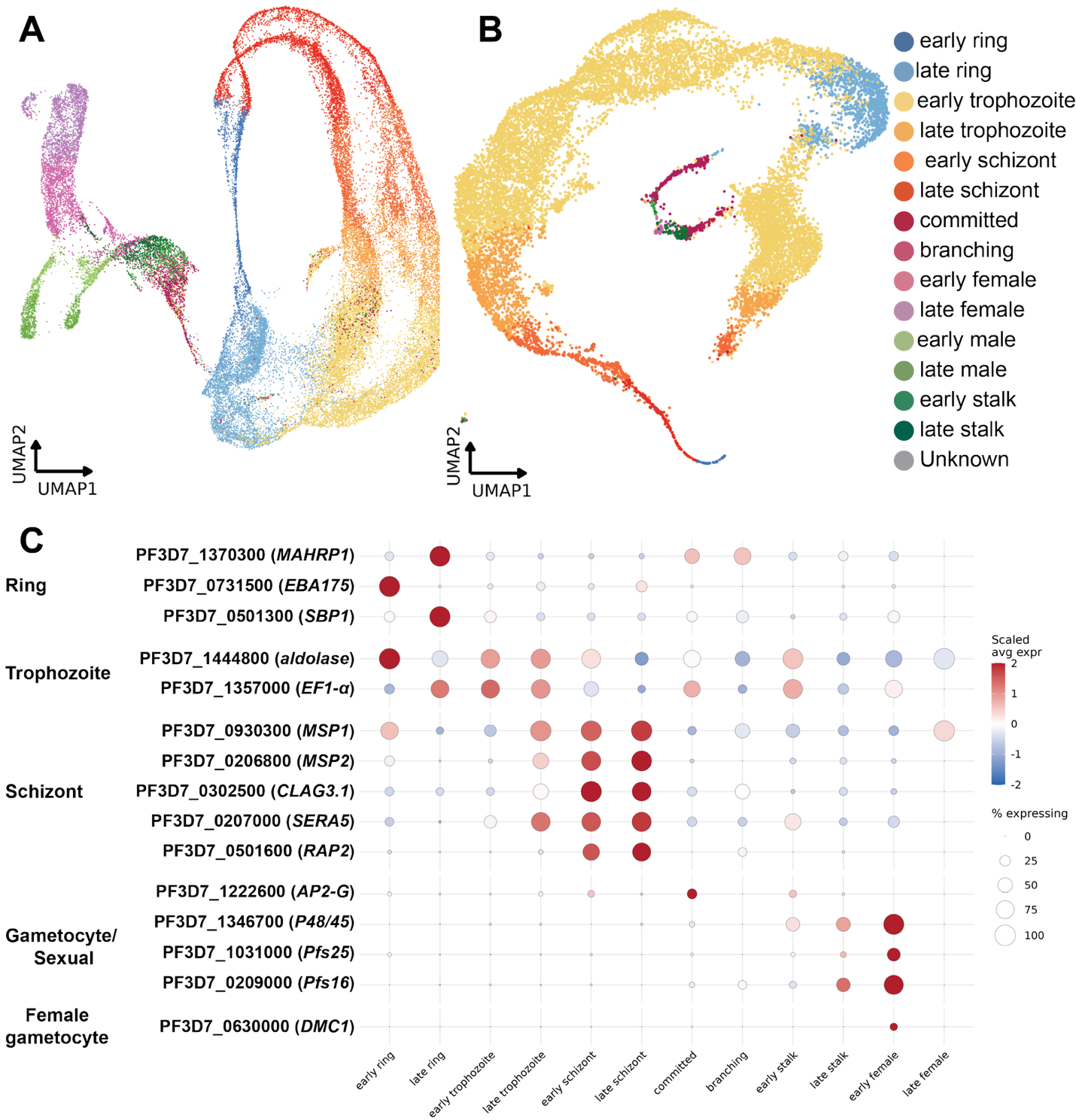

**Supplementary Figure 5. Reference mapping into *P. falciparum* Malaria Cell Atlas recovers full intraerythrocytic developmental cycle confirmed by stage-specific expression marker genes. [A]** UMAP representation of the *P. falciparum* Malaria Cell Atlas (MCA) reference (Dogga et al., 2024), colored by annotated developmental stage, spanning asexual (early/late ring, trophozoite, schizont) and sexual (committed, branching, early/late male and female gametocyte, early/late stalk) populations. **[B]** Transcriptome of captured parasites from the current study projected onto the reference in [A] to assign developmental stage (combined NM and LH conditions). **[C]** Expression of 15 curated stage-specific marker genes across annotated populations in [B], confirming stage-appropriate expression patterns.

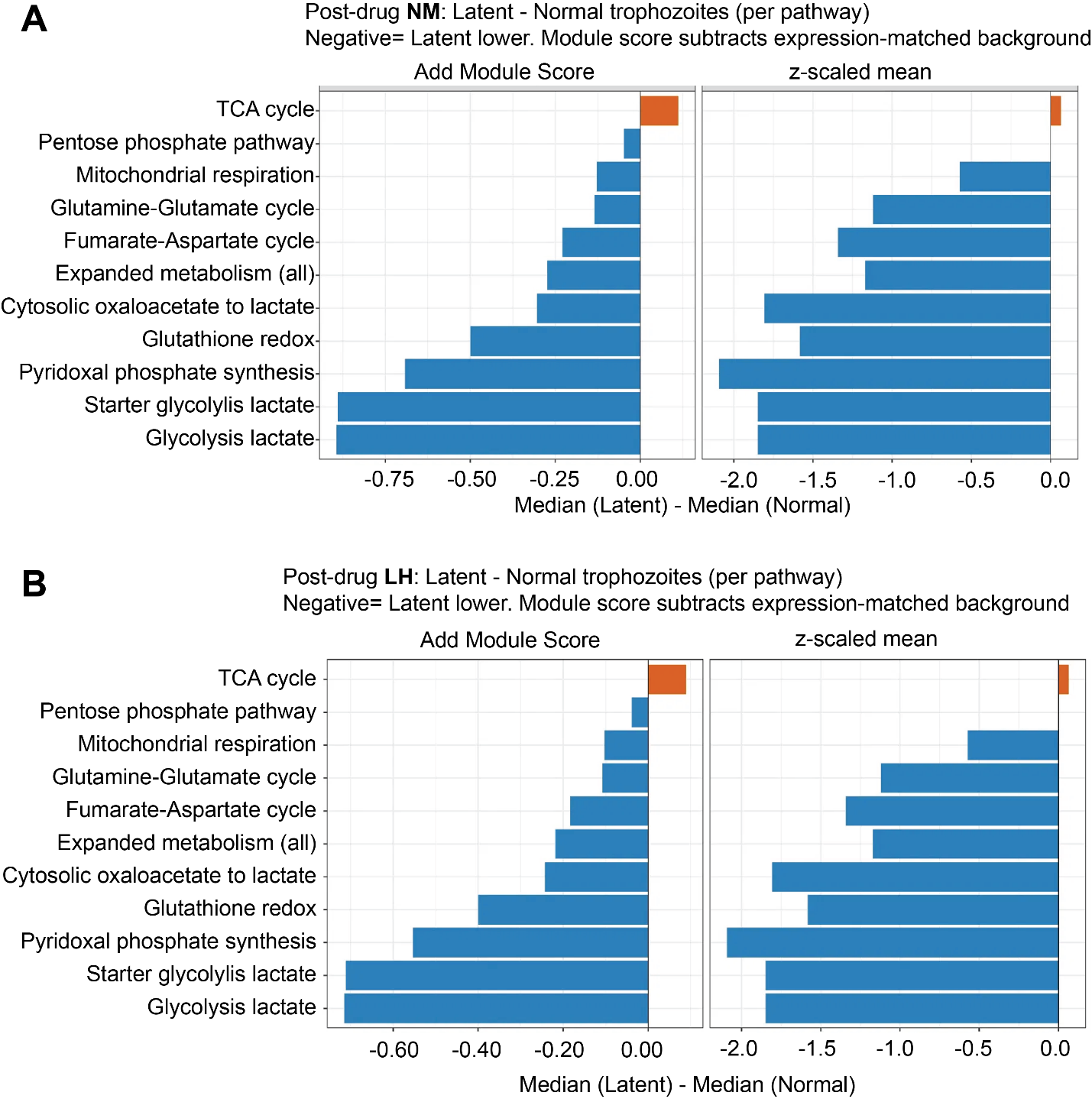

**Supplementary Figure 6. TCA cycle activity is selectively elevated in latent parasites while other metabolic pathway modules are reduced**. For each of 11 metabolic pathway gene sets, we compared per-cell module scores between latent and normal stage-matched parasites in post-DHA [**A**], normal medium (NM) and [**B**] low hypoxanthine (LH) conditions. Scores calculated two ways: Add Module Score (Seurat's AddModuleScore: subtracts an expression-matched background gene set) and a z-scaled mean, shown side by side for each condition. Bars show difference between median latent score and median normal stage-matched score for each pathway (negative values indicate the pathway scores lower in latent than in normal parasites); orange, latent higher than normal; blue, latent lower than normal.
