## Supplementary Table 1, 3-6, 17 for "Environmental stress promotes entry into a pre-existing latent state in *Plasmodium falciparum*"

**Supplementary Table 1. Nutrient deprivation conditions used in this study**.

| **Nutrient Deprivation** | **Class of deprived nutrient** | **Concentration of limited nutrient** | **Length of nutrient deprivation** |
| --- | --- | --- | --- |
| Hypoxanthine | Purine | 0.5µM | 72h |
| Thiamine | Cofactor | 0µM | 48h |

**Supplementary Table 2. Individual experimental replicates underlying nutrient deprivation and latency quantification assays**. Each row represents one independent biological replicate. Date of Experiment: approximate date performed. Line: *P. falciparum* strain used (Dd2). Priming Stress: nutrient-deprivation condition applied prior to drug exposure (low hypoxanthine or thiamine-free). Incubation in medium: duration of nutrient deprivation (72h for hypoxanthine limitation, 48h for thiamine-free conditions). Priming growth reduction %: percent change in parasite growth under nutrient deprivation relative to normal medium control; negative values indicate a growth increase. Non-Primed % Rings and Primed % Rings: percentage of ring-stage parasites in normal medium and nutrient-deprived cultures, confirming comparable staging prior to drug treatment. Priming MMP Change: change in mitochondrial membrane potential under nutrient deprivation relative to control. D10 Drug Stress: DHA concentration used to induce stress and measure parasite recovery up to 10 days post-DHA. D10% Difference: percent difference in growth/parasitemia at day 10 post-drug treatment. Latency Drug Stress: drug concentration used to induce latency (700nM DHA). Synchronization Method: D-Sorbitol or MACS. Latency Proportion in Non-Primed and Latency Proportion in Primed: proportion of parasites entering latency in non-deprived and nutrient-deprived conditions. Fold change in latency: fold change in latency proportion between conditions. D10 growth fold change: fold change in day-10 growth between conditions. eif2aP/Total in Nutrient Deprived and eif2aP/Total in Normal: ratio of phosphorylated to total eIF2α by Western blot.

Supplementary Table 2 (Separate file)

**Supplementary Table 3. Optimization of post-sort recovery medium to select 1% BSA in RPMI for all subsequent sorts.**

| **Exp.**  **no.** | **Sorting buffer^** | **No. cells sorted** | **Pre-sorting^$^** | | **1h post sorting^$^** | | **6h post sorting^$^** | |
| --- | --- | --- | --- | --- | --- | --- | --- | --- |
|  |  |  | **%P** | **%MMP** | **%P** | **%MMP** | **%P** | **%MMP** |
|  | 5% ALB, RPMI | 40,000 | 3.9% | 92% | 76% | 67% | 94% | 91% |
| 1 | RPMI^&^ | 40,000 | 3.9% | 92% | 33% | 98% | 99% | 100% |
|  | **1% BSA, RPMI** | **40,000** | **3.9%** | **92%** | **80%** | **97%** | **90%** | **96%** |
|  | 2% BSA, RPMI | 40,000 | 3.9% | 92% | 91% | 91% | 93% | 96% |
|  | 5% FBS, RPMI | 40,000 | 1.5% | 87% | 65% | 89% | 64% | 86% |
| 2 | 0.04% BSA, RPMI | 40,000 | 1.5% | 87% | 71% | 73% | 61% | 71% |

%P, parasitemia. %MMP, percentage of parasites retaining mitochondrial membrane potential, a measure of viability. ALB, Albumax II; BSA, bovine serum albumin; FBS, fetal bovine serum; **Bold**, media chosen for future sorting steps.

*^*Medium used for sorting and post-sort recovery. For each condition, the same parasite culture was sorted in parallel (40,000 events each) across two different experiments.

**^$^** Timepoints where parasitemia and viability were assessed.

**^&^** Incomplete medium with no serum supplementation.

**Supplementary Table 4. Single-cell RNA sequencing quality metrics pre- and post-DHA treatment.** Cell Ranger-derived sequencing metrics for normal medium (NM) and low hypoxanthine (LH) scRNA-seq libraries, collected before and after DHA treatment.

| **Metric** | **Pre-DHA** | | **Podt-DHA** | |
| --- | --- | --- | --- | --- |
|  | **NM** | **LH** | **NM** | **LH** |
| **Estimated no. of cells** | 12,138 | 9,692 | 5,619 | 8,867 |
| **Mean reads per cell** | 29,449 | 23,113 | 20,024 | 30,465 |
| **Median genes per cell** | 641 | 693 | 463 | 765 |
| **Fraction reads in cells** | 94.60% | 95.60% | 84.50% | 92.70% |

**Supplementary Table 5. Curated stage-marker gene panel used for developmental stage annotation**.

Gene identifiers and products are from PlasmoDB release 68 (*P. falciparum* 3D7); stage assignment reflects the curated marker panel as displayed in the stage-marker dot plot (**Fig. 3B**). "—" indicates no established gene name; PF3D7_0630000 is identified by PlasmoDB ID only.

† EBA-175 (PF3D7_0731500) is canonically a late-schizont/merozoite invasion ligand; it is grouped under Ring in this panel as an invasion/ring-boundary marker.

Curation basis abbreviations: IDC transcriptome, Bozdech et al., PLoS Biol. 2003 (intra-erythrocytic developmental cycle); MCA, Malaria Cell Atlas, Howick et al., Science 2019; sexual atlas, Dogga et al., Science 2024 (single-cell atlas of sexual development); AP2-G commitment, Kafsack et al., Nature 2014 and Sinha et al., Nature 2014.

| **Gene** | **Gene ID** | **Stage marked** | **Product / function** | **Source for curation** |
| --- | --- | --- | --- | --- |
| MAHRP1 | PF3D7_1370300 | Ring | Membrane-associated histidine-rich protein 1; Maurer's-cleft export | Ring/early-trophozoite exported protein. IDC transcriptome (Bozdech et al. 2003); MCA (Howick et al. 2019). |
| EBA-175 | PF3D7_0731500 | Ring † | Erythrocyte-binding antigen-175; merozoite invasion ligand | ring-boundary marker. IDC transcriptome (Bozdech et al. 2003); MCA (Howick et al. 2019). |
| SBP1 | PF3D7_0501300 | Ring | Skeleton-binding protein 1; Maurer's-cleft / knob trafficking | Ring-stage exported protein. IDC transcriptome (Bozdech et al. 2003); MCA (Howick et al. 2019). |
| FBPA | PF3D7_1444800 | Trophozoite | Fructose-bisphosphate aldolase; glycolysis | Trophozoite metabolic peak (glycolytic). IDC transcriptome (Bozdech et al. 2003); MCA (Howick et al. 2019). |
| eEF1α | PF3D7_1357000 | Trophozoite | Elongation factor 1-alpha; translation | Trophozoite peak (high translational activity). IDC transcriptome (Bozdech et al. 2003); MCA (Howick et al. 2019). |
| MSP1 | PF3D7_0930300 | Schizont | Merozoite surface protein 1 | Canonical merozoite/schizont surface antigen. IDC transcriptome (Bozdech et al. 2003); MCA (Howick et al. 2019). |
| RAP2 | PF3D7_0501600 | Schizont | Rhoptry-associated protein 2; invasion organelle | Schizont/merozoite rhoptry protein. IDC transcriptome (Bozdech et al. 2003); MCA (Howick et al. 2019). |
| CLAG3.1 | PF3D7_0302500 | Schizont | Cytoadherence-linked asexual protein 3.1 (clag3) | Schizont-expressed rhoptry/PSAC component. IDC transcriptome (Bozdech et al. 2003); MCA (Howick et al. 2019). |
| MSP4 | PF3D7_0207000 | Schizont | Merozoite surface protein 4 | Merozoite/schizont surface antigen. IDC transcriptome (Bozdech et al. 2003); MCA (Howick et al. 2019). |
| MSP2 | PF3D7_0206800 | Schizont | Merozoite surface protein 2 | Merozoite/schizont surface antigen. IDC transcriptome (Bozdech et al. 2003); MCA (Howick et al. 2019). |
| Pfs48/45 | PF3D7_1346700 | Gametocyte / sexual | 6-cysteine protein P48/45; gamete fertility antigen | Sexual-stage (gametocyte) surface antigen. Sexual atlas (Dogga et al. 2024); MCA (Howick et al. 2019). |
| AP2-G | PF3D7_1222600 | Gametocyte / sexual | ApiAP2 transcription factor AP2-G; master commitment regulator | Master regulator / earliest marker of sexual commitment (Kafsack et al. 2014; Sinha et al. 2014). Sexual atlas (Dogga et al. 2024). |
| Pfs25 | PF3D7_1031000 | Gametocyte / sexual | Ookinete/macrogamete surface protein P25 | Female-biased late sexual / transmission-stage antigen. Sexual atlas (Dogga et al. 2024); MCA (Howick et al. 2019). |
| Pfs230 | PF3D7_0209000 | Gametocyte / sexual | 6-cysteine protein P230; gamete surface / fertility | Sexual-stage (gametocyte) surface antigen. Sexual atlas (Dogga et al. 2024); MCA (Howick et al. 2019). |
| — | PF3D7_0630000 | Female gametocyte | CPW-WPC family protein | Female-gametocyte-enriched CPW-WPC family gene. Sexual atlas (Dogga et al. 2024). |

**Supplementary Table 6. Gene-set and module definitions**

**Stage validation - curated canonical markers**

| **Stage/module** | **Genes** |
| --- | --- |
| Ring | MAHRP1 (PF3D7_1370300), EBA175 (PF3D7_0731500), SBP1 (PF3D7_0501300) |
| Trophozoite | Aldolase (PF3D7_1444800), EF1-α (PF3D7_1357000) |
| Schizont | MSP1 (PF3D7_0930300), MSP2 (PF3D7_0206800), CLAG3.1 (PF3D7_0302500), SERA5 (PF3D7_0207000), RAP2 (PF3D7_0501600) |
| Gametocyte/sexual | AP2-G (PF3D7_1222600), P48/45 (PF3D7_1346700), Pfs25 (PF3D7_1031000), Pfs16 (PF3D7_0209000) |
| Female gametocyte | DMC1 (PF3D7_0630000) |

**Stage validation - expanded marker set**

| **Stage/module** | **Genes** |
| --- | --- |
| Ring | SBP1 (PF3D7_0501300), KAHRP (PF3D7_0202000), EBA175 (PF3D7_0731500), PfEMP3 (PF3D7_1300300), SMS1 (PF3D7_0424400), MAHRP1 (PF3D7_1370300) |
| Trophozoite | CRT (PF3D7_0709000), MDR1 (PF3D7_0523000), EF1-α (PF3D7_1357000), aldolase (PF3D7_1444800), HSP70 (PF3D7_0818900) |
| Schizont | MSP1 (PF3D7_0930300), MSP2 (PF3D7_0206800), AMA1 (PF3D7_1133400), CLAG3.1 (PF3D7_0302500), SERA5 (PF3D7_0207000), RAP2 (PF3D7_0501600) |
| Gametocyte/sexual | AP2-G (PF3D7_1222600), GDV1 (PF3D7_0935400), P48/45 (PF3D7_1346700), Pfs25 (PF3D7_1031000), Pfs16 (PF3D7_0209000), Pfs230 (PF3D7_0408600) |
| Female gametocyte | GAP27/25 (PF3D7_1302100), DMC1 (PF3D7_0630000) |
| Male gametocyte | Dynein (PF3D7_1438800), MD1 (PF3D7_1214800), HAP2 (PF3D7_1014200) |
| **S**exual commitment | AP2-G (PF3D7_1222600), GDV1 (PF3D7_0935400), P48/45 (PF3D7_1346700), Pfs25 (PF3D7_1031000), Pfs16 (PF3D7_0209000), Pfs230 (PF3D7_0408600), DMC1 (PF3D7_0630000). |

**Sexual-commitment**

Cells were classified as sexual-marker-high if their sexual-marker score exceeded the 95th percentile of the normal early trophozoite sexual score.

**Data-driven stage signatures**

Data-driven stage signatures were derived across the following predicted stage categories: early ring, late ring, early trophozoite, late trophozoite, early schizont, late schizont, committed, branching, early stalk, late stalk, early female, late female, early male, and late male. The normal early trophozoite signature was derived from normal early trophozoites only.

**Metabolic modules**

| **Module** | **Genes** |
| --- | --- |
| Starter glycolysis/lactate | Hexokinase (PF3D7_0624000), aldolase (PF3D7_1444800), PGK (PF3D7_0922500), GAPDH (PF3D7_1462800), enolase (PF3D7_1015900), pyruvate kinase (PF3D7_0626800), LDH (PF3D7_1324900) |
| TCA cycle | Citrate synthase (PF3D7_1022500), aconitase (PF3D7_1342100), isocitrate dehydrogenase (PF3D7_1345700), α-ketoglutarate dehydrogenase components (PF3D7_1312600, PF3D7_0504600), succinyl-CoA synthetase components (PF3D7_1108500, PF3D7_1431600), succinate dehydrogenase components (PF3D7_1212800, PF3D7_1034400), fumarate hydratase/FH (PF3D7_0927300), malate:quinone oxidoreductase/MQO (PF3D7_0616800) |
| Fumarate/aspartate cycle | Aspartate aminotransferase (PF3D7_0204500), malate dehydrogenase (PF3D7_0618500), adenylosuccinate synthetase (PF3D7_1354500), adenylosuccinate lyase (PF3D7_0206700), AMP deaminase (PF3D7_1329400) |
| Cytosolic oxaloacetate-to-lactate | PEPCK (PF3D7_1342800), PEPC (PF3D7_1426700), pyruvate kinase (PF3D7_0626800), LDH (PF3D7_1324900) |
| Mitochondrial respiration | G3PDH (PF3D7_1114800), NDH2 (PF3D7_0915000), DHODH (PF3D7_0603300), complex III components (PF3D7_1439400, PF3D7_1462700, PF3D7_1012300), complex IV components (PF3D7_0927800, PF3D7_0928000, PF3D7_1475300), complex V/ATP synthase components (PF3D7_1235700, PF3D7_0217100, PF3D7_0715500), cytochrome c (PF3D7_1404100) |
| Glutamine/glutamate | Glutamate synthase (PF3D7_1435300), glutamate dehydrogenases (PF3D7_0802000, PF3D7_1416500, PF3D7_1430700) |
| Glutathione/redox | Glutathione synthetase (PF3D7_0512200), glutathione reductase (PF3D7_1419800), γ-glutamylcysteine synthetase/γ-GCS (PF3D7_0918900), glutaredoxin (PF3D7_0306300), thioredoxin (PF3D7_1457200) |
| Pentose phosphate pathway | G6PD-6PGL/GluPho (PF3D7_1453800), 6-phosphogluconate dehydrogenase/6PGD (PF3D7_1454700) |
| Pyridoxal phosphate synthesis | PDX2 (PF3D7_1116200), PDX1 (PF3D7_0621200), pyridoxal kinase (PF3D7_0616000) |

**Supplementary Table 7. MAST DE Analysis of all genes Latent vs Normal (Separate file)**

**Supplementary Table 8. Latent vs Normal only Up (Separate file)**

**Supplementary Table 9. Latent vs Normal only down (Separate file)**

**Supplementary Table 10. Latent vs Normal GSEA (Separate file)**

**Supplementary Table 11. Latent vs Normal Absolute GSEA (Separate file)**

**Supplementary Table 12. Mixed Pathways Latent vs Normal (Separate file)**

**Supplementary Table 13. GO Enrichment Latent vs Normal (Separate file)**

**Supplementary Table 14. Differential expression and blood-stage essentiality of metabolic genes represented in Figure 6 (Separate file)**

**Supplementary Table 15. Differential expression and blood-stage essentiality of stress-response and proteostasis genes in latent versus normal stage matched parasites (Separate file)**

**Supplementary Table 16. Curated antigen trafficking and variant surface antigen gene families, with differential expression status in latent versus normal parasites (Separate file)**

**Supplemental Table 17. Uncharacterized differentially expressed genes and their assignment within the latency program.** Summary of PlasmoFP-based functional assignment among genes lacking prior functional annotation (proteins of unknown function) that were differentially expressed between latent and normal parasites (latent-versus-normal MAST analysis). Of 1,388 total uncharacterized DE genes (902 upregulated, 486 downregulated in latent), 809 (58%) received at least one PlasmoFP Gene Ontology (GO) prediction at FDR ≤ 0.05: 527 (38%) were newly annotatable (no prior GO annotation), and 282 (20%) had a prior GO annotation corroborated by the PlasmoFP prediction. Within the significant subset (adjusted P < 0.05, |log₂FC| ≥ 1; 675 genes: 495 upregulated, 180 downregulated), 406 (60%) received a PlasmoFP prediction, of which 275 (41%) were newly annotatable. Predicted functional themes are shown for the 406 annotated genes in this significant subset, with percentages calculated as the proportion of those 406 genes bearing a predicted term in each theme; themes are not mutually exclusive, so a single gene may be counted under more than one theme and percentages do not sum to 100%.

| **Category** | **Genes (n)** | **% of total** |
| --- | --- | --- |
| **All uncharacterized differentially expressed genes** |  |  |
| Total uncharacterized DE genes | 1,388 | 100% |
| Upregulated in latent | 902 | 65% |
| Downregulated in latent | 486 | 35% |
| Received ≥1 PlasmoFP GO prediction (FDR ≤ 0.05) | 809 | 58% |
| newly annotatable (no prior GO annotation) | 527 | 38% |
| had prior GO (corroborating prediction) | 282 | 20% |
| **Significant subset (adj. P < 0.05 and \|log₂FC\| ≥ 1)** |  |  |
| Significant uncharacterized genes | 675 | 100% |
| Upregulated / downregulated | 495 / 180 | 73% / 27% |
| Received ≥1 PlasmoFP prediction | 406 | 60% |
| newly annotatable | 275 | 41% |
| **Predicted functional themes (significant subset)ᵃ** |  |  |
| ***Theme*** | ***Genes with ≥1 term*** | ***% of annotatedᵇ*** |
| Binding / catalytic activity | 303 | 75% |
| Membrane / organellar localization | 45 | 11% |
| Nucleotide / small-molecule binding | 44 | 11% |
| Regulation of macromolecule / RNA biosynthesis | 39 | 10% |
| Hydrolase / transferase activity | 31 | 8% |
| Structural constituent of ribosome / translation | 29 | 7% |
| Cytoskeletal / tubulin binding | 19 | 5% |
| RNA processing / binding | 13 | 3% |
| Transmembrane transport | 10 | 2% |
| Oxidoreductase / redox | 5 | 1% |
| Kinase / phosphatase signaling | 4 | 1% |

**Supplementary Table 18. MAST DE Analysis of all genes LH latent vs NM Latent (Separate file)**

**Supplementary Table 19. LH latent vs NM Latent only Up (Separate file)**

**Supplementary Table 20. LH latent vs NM Latent only down (Separate file)**

**Supplementary Table 21. LH latent vs NM Latent GSEA (Separate file)**

**Supplementary Table 22. LH latent vs NM Latent Absolute GSEA (Separate file)**

**Supplementary Table 23. Mixed Pathways LH latent vs NM Latent GO (Separate file)**

**Supplementary Table 24. Separate f200 gene classifier (Separate file)**
